# MLG: Multilayer graph clustering for multi-condition scRNA-seq data

**DOI:** 10.1101/2021.03.26.437231

**Authors:** Shan Lu, Daniel J. Conn, Shuyang Chen, Kirby D. Johnson, Emery H. Bresnick, Sündüz Keleş

**Affiliations:** Department of Statistics, University of Wisconsin, Madison, WI, USA; Department of Biostatistics and Medical Informatics, University of Wisconsin School of Medicine and Public Health, Madison, WI, USA; Wisconsin Blood Cancer Research Institute, Department of Cell and Regenerative Biology, University of Wisconsin School of Medicine and Public Health, Madison, WI, USA

**Keywords:** single cell RNA-sequencing, dimension reduction, clustering, stochastic block models, data integration

## Abstract

Single-cell transcriptome sequencing (scRNA-seq) enabled investigations of cellular heterogeneity at exceedingly higher resolutions. Identification of novel cell types or transient developmental stages across multiple experimental conditions is one of its key applications. Linear and non-linear dimensionality reduction for data integration became a foundational tool in inference from scRNA-seq data. We present **M**ulti **L**ayer **G**raph Clustering (MLG) as an integrative approach for combining multiple dimensionality reduction of multi-condition scRNA-seq data. MLG generates a multilayer shared nearest neighbor cell graph with higher signal-to-noise ratio and outperforms current best practices in terms of clustering accuracy across large-scale bench-marking experiments. Application of MLG to a wide variety of datasets from multiple conditions highlights how MLG boosts signal-to-noise ratio for fine-grained sub-population identification. MLG is widely applicable to settings with single cell data integration via dimension reduction.

## 1 Introduction

High-throughput single-cell RNA sequencing (scRNA-seq) captures transcriptomes at the individual cell level. While scRNA-seq is powerful for a wide range of biological inference problems, perhaps, its most common application thus far is cell type/stage identification. Specifically, clustering analysis of scRNA-seq enables identification of cell types in tissues [1–3], or discrete stages in cell differentiation and development [4, 5] by leveraging similarities in the transcriptomes of cells. Although a plethora of methods, some of which repurpose existing k-means and Louvain algorithms for clustering, exist for cell type identification from single conditions [6], clustering of scRNA-seq data from multiple biological conditions (e.g., different treatments, time points, tissues) to elucidate cell types and subpopulation of cells has not been a major focus.

Joint clustering of scRNA-seq datasets across multiple conditions entails, in addition to standard normalization, feature selection, and dimension reduction, a consideration of whether or not data from cells across multiple stimuli need to be “integrated” before the downstream analysis of clustering. This is because cell type-specific response to experimental conditions may challenge a joint analysis by separating cells both by experimental condition and cell type. Most notably, Seurat (v3) [7] uses canonical component analysis (CCA) [8] to perform data integration for high dimensional gene expression across conditions. Liger [9] achieves dimension reduction and data integration simultaneously by using penalized nonnegative matrix factorization (NMF) to estimate factors shared across conditions and specific to conditions. ScVI [10] and scAlign [11] both use deep neural networks to infer a shared nonlinear low-dimensional embedding for gene expression across conditions. Harmony [12] iteratively performs soft k-means clustering and condition effect removal based on clustering assignments. A common theme of these approaches is that majority of them (Liger, scAlign, scVI, and Harmony) directly yield low-dimensional integrated data for downstream visualization and clustering.

While existing methods for joint analysis of scRNA-seq data (e.g., Seurat, scAlign, and Liger among others) regularly adapt modularity maximization algoritms such as Louvain graph clustering [13] with shared nearest neighbor (SNN) graph or shared factor neighborhood (SFN) graph built with their low-dimensional embeddings of the data, they vastly differ in their dimension reduction techniques as we outlined above. This leads to documented notable differences among these methods [14]. To leverage strengths of different dimensionality reduction approaches, we develop an integrative framework named multilayer graph (MLG) clustering as a general approach that borrows strength among a variety of dimension reduction methods instead of focusing on a single one that best preserves the cell-type specific signal. MLG takes as input a set of low-dimensional embeddings of all the cells, integrates them into a shared nearest neighbour graph with analytically provable improved signal-to-noise ratio, and clusters them with the Louvain algorithm, which has recorded outstanding performances in benchmarks [15, 16] and is also the default clustering algorithm in many scRNA-seq analysis packages such as Scanpy [17], Seurat (v2, v3) [7, 18] and Liger [9]. MLG framework leverages a key observation that the nearest neighbor graphs constructed from different dimensionality reductions of scRNA-seq data tend to have low dependence. A consequence of this observation, supported by analytical calculations, is that the resulting multilayer integrative scheme yields a combined cell graph with higher signal-to-noise ratio. We further corroborate this result with computational experiments using benchmark data and illustrate that MLG clustering outperforms current best practices for jointly clustering cells from multiple stimuli and preserves salient structures of scRNA-seq data from multiple conditions. We illustrate this property of MLG clustering with an application to scRNA-seq data from mouse hematopoietic stem and progenitor cells (HSPCs) under two conditions (with or without a *Gata2* enhancer deletion)[19], and from mouse HSPCs under four conditions (*Gif1* ^+*/*+^, *Gif1* ^*R*412*X/R*412*X*^, *Gif1* ^*R*412*X/−*^, *Gif1* ^*R*412*X/−*^ *Irf8* ^+*/−*^) [20]. Finally, we showcase how MLG enables robust analysis of recent SNARE-seq [21] data which generates two data modalities, accessible chromatin and RNA, within the same cells.

## 2 MATERIALS AND METHODS

### 2.1 Overview of Multilayer Graph Clustering (MLG)

The MLG clustering algorithm is a general framework that aggregates shared nearest neighbourhood graphs constructed from different linear and non-linear low-dimensional embeddings of scRNA-seq data (Figure 1A). It takes as input *G* sets of low-dimensional embeddings of the same dataset generated by different dimensionality reduction methods, with and/or without data integration for datasets across multiple conditions. Consequently, it constructs and then aggregates SNNs from each of the *G* embeddings and leverages Louvain modularity [13] maximization algorithm for the final clustering. Figure 1A depicts the workflow of the MLG with four existing scRNA-seq low-dimensional embedding methods as inputs. Here, we considered dimension reduction with PCA and consensus nonnegative matrix factorization (cNMF) [22] as representatives of low-dimensional embeddings without data integration across multiple conditions, and Seurat [7] and Liger [9] as representatives with data integration. Next, we describe the construction of the SNN graphs in detail.

**Figure 1.**
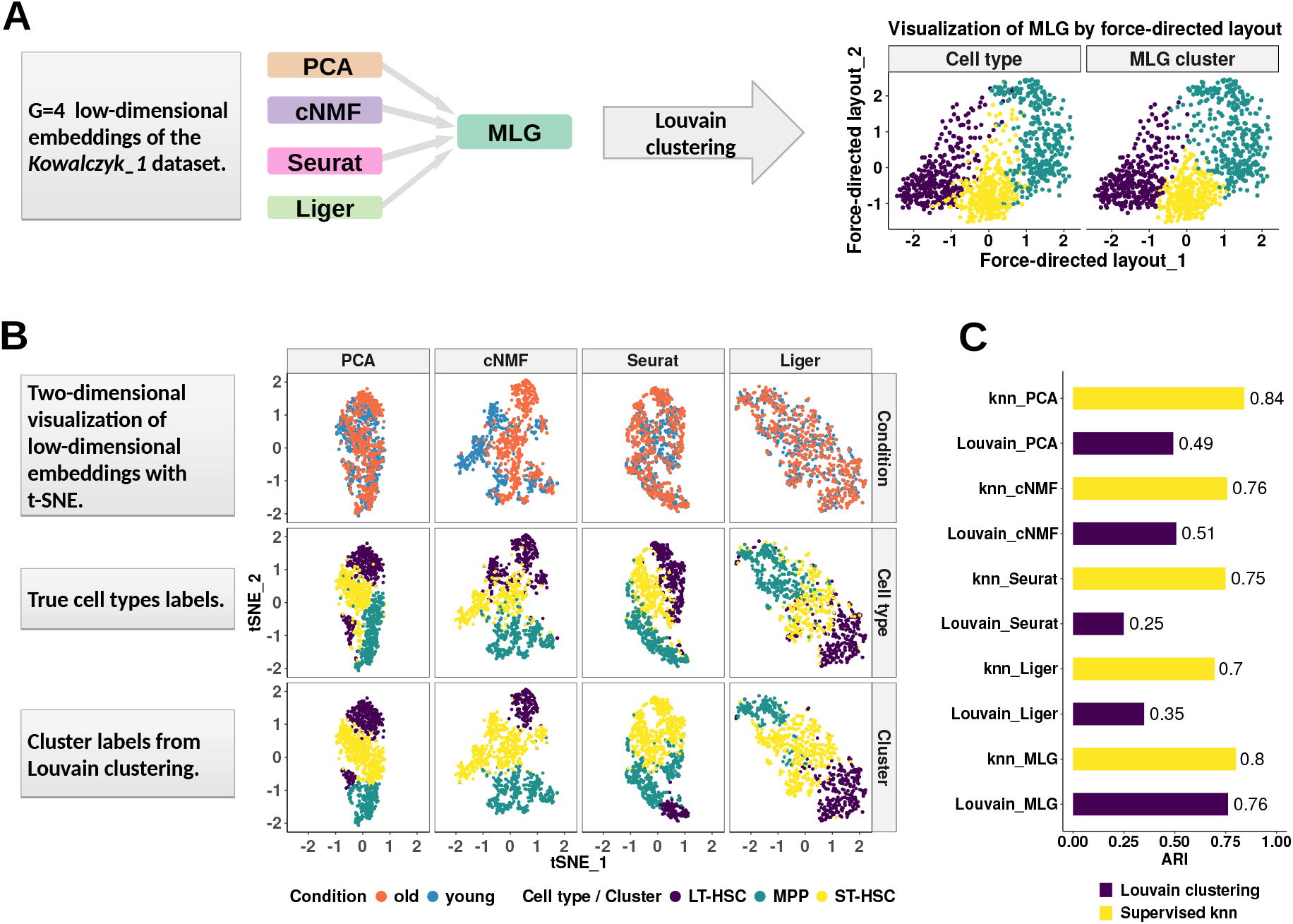
Multilayer graph (MLG) clustering workflow with an illustration on the *Kowalczyk 1* dataset. (A) MLG takes as input G different low-dimensional embeddings to construct SNN graphs, aggregates the resulting graphs, and applies Louvain algorithm for clustering the aggregated graph. The two scatter plots on the right visualize the MLG clustering results with a force-directed layout where the cells are labeled with their true labels (left) and MLG clustering labels (right). (B) Visualization of the PCA, cNMF, Seurat integration and Liger factors with t-SNE coordinates. Cells are labeled according to underlying experimental conditions (top row), cell types (middle row), and Louvain clustering assignments from SNN graphs constructed with each dimension reduction method (bottom row). (C) Adjusted Rand Index (ARI) between true cell type labels and labels from clustering/knn classification.

Let *D ∈* ℝ^*n×d*0^, where *n* and *d*_0_ represent the number of cells and the dimension of latent factors, respectively, denote a low-dimensional embedding matrix. As in Figure 1A, the matrix *D* can be obtained from principle component analysis (PCA), consensus nonnegative matrix factorization (cNMF) [22] or low-dimensional embeddings from scRNA-seq analysis packages such as scVI, Liger applied to scRNA-seq count matrices. For each cell *i*, let *L*_*k*_(*i*) denote the set of *k* nearest neighbours based on Euclidean distance across the *d*_0_ latent factor vectors. An edge is added to the undirected SNN graph (i.e., *A*_*ij*_ is set to 1 in the corresponding adjacency matrix *A*) between cells *i* and *j* if their number of common neighbors is larger than *kλ*, where *λ* is a filtering threshold. The parameters *k* and *λ* determine the sparsity level of the SNN graph. A large *k* and small *λ* lead to a denser graph. While there is no optimal criteria to pick these parameters, Von Luxburg [23] suggests *k* = *O*(*n*) to guarantee a connected graph. In the analyses provided in this paper, we used *k* = 20 and *λ* = 1*/*5. We also showed with our simulation study that clustering accuracy is relatively robust to the choice of *k*.

Next, given *G* adjacency matrices *{A*^(*g*)^*}, g* = 1,*…, G*, each corresponding to an SNN from a low-dimensional embedding, we aggregate them into a union adjacency matrix *B* as follows:

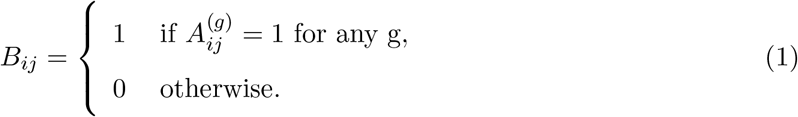

MLG then uses this resulting adjacency matrix *B* for clustering with the Louvain modularity [13] maximization algorithm.

### 2.2 Benchmark datasets

We leveraged three public scRNA-seq datasets, summarized as (log-normalized) counts and with complementary characteristics, to benchmark MLG (Supplementary Table S1). The first dataset, that we refer to as *Kowalczyk 1*, profiled transcriptomes of HSPCs among mice of different ages (*young* at 2-3 months and *old* at 22 months) and harbored cells from three hematopoietic stages: long-term (LT)-HSC, short-term (ST)-HSC, and multi-potent progenitor (MPP) [24]. We used this dataset for explicitly illustrating how different dimension reduction algorithms operate on scRNA-seq data across multiple conditions. An extended version of this dataset, labelled as *Kowalczyk 2*, included additional LT-HSC and ST-HSC cells from independent mice that were profiled months apart from the original dataset. A third dataset from the HSPC system [25] included the same cell types (LT-HSC, ST-HCS, and MPP) from young and old mice and with and without LPS+PAM stimuli. This dataset (referred to as *Mann*) is an example with two factors at two levels and represents a setting with four experimental conditions. The third dataset, labelled as *Cellbench* [26], included scRNA-seq data of 636 synthetic cells created from three human lung adenocarcinoma cell lines HCC827, H1975, and H2228. RNA was extracted in bulk for each cell line. Then each cell line’s RNA was mixed in at seven different proportions and diluted to single cell equivalent amounts ranging from 3.75pg to 30pg.

### 2.3 Aggregation of multiple SNNs boosts signal-to-noise ratio

Next, we define a notion of signal-to-noise ratio for graphs and show that MLG aggregation of SNNs boosts the signal-to-noise ratio of the sparse graphs typically found in scRNA-seq analyses. We base our theoretical analysis on stochastic block models (SBMs) [27], which are generative random graph models that serve as canonical models for investigating graph clustering and community detection. In the view of SBMs, cells are represented as vertices in a binary graph with edges representing similarity between cells. The cells are assumed to reside in distinct communities determined by cell-type, and this community structure determines the probability of an edge between cells. We denote the total number of cells (vertices) by *n* and the number of communities (clusters) by *K*. Let *σ* be a cluster assignment function where *σ*(*i*) corresponds to the cluster label for cell *i*, and *θ*_*σ*(*i*)*σ*(*j*)_ denotes the connectivity probability (i.e., edge probability) between cells *i* and *j*. Furthermore, let *θ*_*in*_ and *θ*_*out*_ denote the minimum in-cluster and maximum out-of-cluster connectivity probability. We define the signal-to-noise ratio of the SBM as

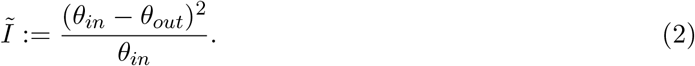

Intuitively, if the difference between the maximum in-cluster and out-of-cluster connectivity probabilities is large, it will be easier to distinguish communities from one another. Furthermore, this intuition is supported by theory, as Zhang et al. [28] prove that the performance of an optimal SBM estimator depends heavily on *Ĩ*. We provide a rigorous explication of this theoretical result in the Supplementary Materials.

We utilize this specific notion of signal-to-noise ratio to investigate the impact of aggregation. Given adjacency matrices *A*_1_ and *A*_2_ of two independent SBM graphs on the same set of cells, their union adjacency matrix *B*, we use superscripts to indicate parameters of each graph, e.g., 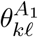 represents the connectivity probability of cells in cluster *k* and cells in cluster *𝓁* in graph *A*_1_. In the realm of scRNA-seq data *A*_1_ and *A*_2_ are usually quite sparse, i.e., with small 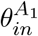 and 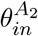. If we further assume that 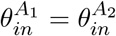 and 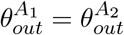, we arrive at the primary result of this section

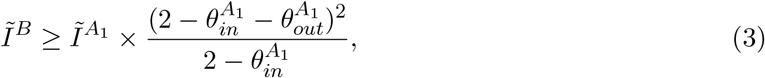

which implies that the signal-to-noise ratio of the aggregated SBM is nearly twice that of either of the base graphs, *A*_1_ and *A*_2_. We provide a detailed derivation of this result in the Supplementary Materials.

This result on improved signal-to-noise ratio due to aggregation is foundational for establishing operational characteristics of MLG. It assumes that the SBMs are both sparse and independent of one another. As we show in the Results section, empirical observation also supports this assumption as SNN graphs are sparse, and we see only small proportions of overlapping edges between SNN graphs of different low-dimensional projections. In the Supplementary Materials, we further show that dimension reduction leads to perturbation in the local neighborhood structure of the full-data (e.g., data prior to dimension reduction) SNN graph. This, in turn, indicates that the dependence between SBMs derived from different low-dimensional projections is weak, and thus the overall assumptions of our result on the benefits of aggregation are approximately met. This result also explains the empirical behavior concerning low proportions of overlapping edges between SNN graphs that we observe across our benchmarking datasets in the Results section.

#### Signal-to-noise ratio estimation

For benchmark datasets with true cell type labels, an empirical version of signal-to-noise ratio for a given cell graph can be calculated as:

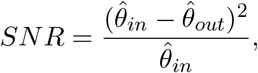

where 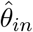 stands for the estimated minimum “within cell type” connectivity probability and 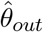 stands for the estimated maximum “out-of-cell type” connectivity probability. Let *A ∈ {*0, 1*}*^*n×n*^ denote the adjacency matrix of a cell graph with *n* cells from *K* cell types and let *C*_*k*_ denote the set of cells in cell type *k*. Then, 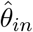 and 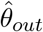 are given by

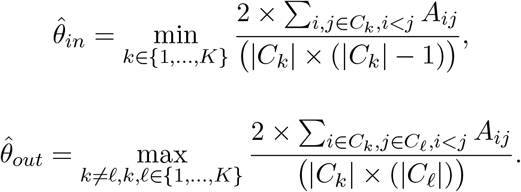

#### Details on the execution of different scRNA-seq analysis methods

##### PCA

PCA is implemented through `Seurat (version 3.1.4)` package. Specifically, we used Seurat function `NormalizeData` to scale counts with a size factor of 10,000 and then performed log-transformation while adding a count of 1 to avoid taking log of 0. The functions `FindVariableFeatures` and `ScaleData` were used to find highly variable features and adjust for technical and biological effects, respectively. The condition labels and total genes counts were regressed out with `ScaleData` for all simulated, benchmark, and application datasets. The percentage of mitochondrial gene counts were also regressed out for the dataset *Johnson20*. Each gene is scaled by its standard error of counts across cells before performing PCA with the `RunPCA` function.

##### cNMF

cNMF is applied through the code provided by [22] on Github https://github.com/dylkot/cNMF. Count matrices were provided as input for cNMF and normalization and feature selection were carried out within the cNMF pipeline. The consensus analysis used 50 NMF runs.

##### Seurat-integration

Following the Seurat tutorial, function `SplitObject` was used to split dataset by conditions, then `FindIntegrationAnchors` and `IntegrateData` were applied to integrate gene expression across conditions. The integrated gene expression matrix was scaled with function `ScaleData`. *RunPCA* was used to reduce dimension for the scaled and integrated gene expression matrix.

##### Liger

Gene expression count matrices were normalized with the R package `Liger(0.4.2)` function `normalize`. Highly variable genes were chosen with `selectGenes`. Functions `scaleNotCenter, optimizeALS` and `quantileAlignSNF` were utilized to scale genes expression, implement data integration and Liger default clustering, respectively. We used the low-dimension embedding in data slot @H by scaling it to have row sum of 1 in SNN graph construction for MLG.

##### scAlign

Our application followed the tutorial of scAlign on Github https://github.com/quon-titative-biology/scAlign. To reduce computation time, we use CCA factors as input for the encoder neural network for dataset *Kowalczyk 1* and *Kowalczyk 2*. Dataset *Mann* has 4 conditions, but multi-CCA are not implemented in scAlign. We use PCA factors as input for *Mann*.

##### Harmony

We applied `Harmony` through the *Seurat* workflow with the `RunHarmony` function. nclust parameter was set to the true numbers of clusters for both the simulated and benchmark datasets.

##### scVI

ScVI is implemented through the scVI python package, available at https://scvi.readthedocs.io/en/stable/. We followed its “basic tutorial” for model training, and used sample latent factors as input for k-means or Louvain clustering.

##### Parameters common to all methods

We kept 3,000 genes for all the benchmark scRNA-seq datasets and 2,500 genes for the simulated datasets. We kept 15 latent factors from all dimension reduction or data integration methods for the analysis of the benchmark scRNA-seq datasets. We applied k-means with the R function kmeans and set the numbers of clusters to the true number of clusters all the datasets. Louvain clustering was applied through `Seurat::FindClusters`. The resolution was chosen through grid search until the target number of clusters was met. The target number of clusters were set to the true numbers of clusters for the benchmark datasets. The numbers of clusters for *Johnson20* and *Muench20* was chosen by the eigengap heuristic [23].

##### Differential expression analysis

Differential expression analysis was carried out with function `Seurat::FindMarkers` using the “MAST” algorithm. In the case of multiple conditions per dataset, cluster markers were identified with a differential expression analysis of cells from the same condition across different clusters and the p-value was set to be the smallest among all conditions.

The precision-recall values are based on gold standard cell type marker genes and cell type-specific DE genes across conditions, defined as having Bonferroni corrected p-values less than 0.01 in the analysis with the true cell labels. The precision-recall values were reported at cutoffs of 0.2, 0.1, 0.05, 0.01, 0.001 for Bonferroni adjusted p-values in the cell type marker gene and condition DE gene identification analysis.

##### Gene set enrichment analysis

Gene set enrichment analysis was carried out with the R package `topGO` (2.36.0) using the Fisher’s exact test and elim [29] algorithm.

##### Processing of the SNARE-seq data

SNARE-seq generates joint profiles of accessible chromatin (snATAC-seq) and RNA (snRNA-seq). Peak matrices and expression matrices for the first replication of neonatal and adult mouse brain cortex provided along with the raw data are used in this analysis. Fragment files for integration are extracted from the raw snATAC-seq data using the python package Sinto. Following the Seurat tutorial, functions `FindIntegrationAnchors` and `IntegrateData` were applied to peak matrices with fragment files to integrate chromatin accessibility data across conditions. We used Latent Semantic Indexing [30] (LSI) as the dimension reduction method for snATAC-seq data and removed the first LSI component from downstream analysis since it often captures sequencing depth rather than biological variation. SnRNA-seq data are processed with Seurat and Liger integration. Weighted Nearest Neighbor [31] analysis was then applied to different combinations of dimension reduction results (LSI-Seurat, LSI-Liger).

##### Simulations

Our simulation set-up is based on an adaptation of the simulation setting of [22] to allow differential expression across conditions and takes advantage of commonly used scRNA-seq simulation models from Splatter [32]. In this set-up, the mean gene-expression profile for each cell is a weighted sum of *cell identity gene expression program* (GEP) and *activities* GEPs. Here, the cell identity GEP characterizes cell types whereas the activity GEP represents other sources of biological variations, like cell cycle effects or responses to specific stimuli. This set up also mimics the datasets with multiple cell types under stimuli (e.g., HSPCs from old and young mouse), that we have utilized in this paper. Specifically, each simulation replication involved the following data generation process.

1. **Simulate base mean expression**. For gene *w* (*w* = 1,…, *W*), sample the magnitude of mean expression 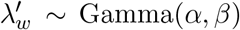, the outlier indicator 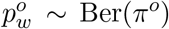, and the outlier factor 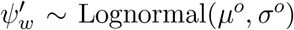. The base mean expression for gene *w* is defined as 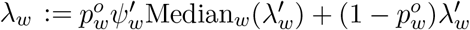
2. **Simulate gene expression program (GEP)**. For gene *w*, sample differential expression (DE) indicator *p*_*w*_ *∼* Ber(*π*), DE factor *ψ*_*w*_ *∼* Lognormal(*µ, σ*), and down regulation indicator 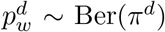.Let 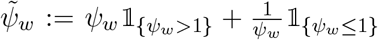. The DE ratio for gene *w* is defined as 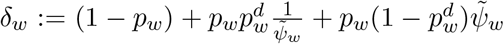. The GEP is given by 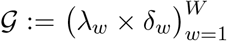, where *W* is the total number of genes. Here, *λ*_*w*_ and *δ*_*w*_ denote the base gene mean and DE ratio of gene *w*, respectively. This procedure can be used to simulate identity and activity GEPs, with slightly different set of parameters (*π, π*^*d*^, *µ, σ*).
3. **Simulate mean gene expression for cell type** *t* **and condition** *c* (*t* = 1,…, *T, c* = 1*… C*). Following the procedure in step 2, simulate identity GEP 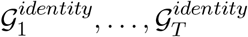 and activity GEP 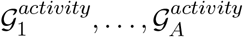. Given activity GEP weights 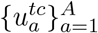 for cell type *t* and condition *c*, the cell type and condition specific mean gene expression is defined as 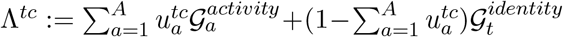. Activity GEP weights control the magnitude of condition effects.
4. **Correct for library size**. For cell *k* in cell type *t* and condition *c*, sample the library size for cell *k L*_*k*_ *∼* Lognormal(*µ*^*l*^, *σ*^*l*^). The mean gene expression of cell *k* after correction for the library size is 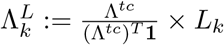.
5. **Correct for biological coefficient of variation (BCV)**. Let *φ* be the BCV dispersion parameter, `df` be the degrees of freedom of the BCV inverse *χ*^2^ distribution. The BCV for gene *w* of cell *k* is sampled through the formula 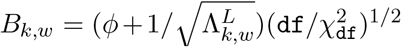. The BCV corrected mean for gene *w* of cell *k* is sampled by 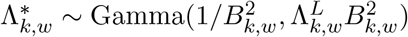
6. **Generate counts**. Sample counts for gene *w* of cell 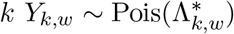.

In order to make these simulations realistic, we set the parameters in a data-driven way. Specifically, the parameters of the simulation setting (*α* = 1.46, *β* = 1.48, *π*^*o*^ = 0.091, *µ*^*o*^ = 2.82, *σ*^*o*^ = 0.84, *µ*^*l*^ = 8.95, *σ*^*l*^ = 0.45, *φ* = 0.11, df = 36.57) are estimated from the *Johnson20* dataset with the Splatter package. Parameters generating identity and activity GEPs are (*π* = 0.025, *π*^*d*^ = 0.2, *µ* = 0.65, *σ* = 0.25) and (*π* = 0.05, *π*^*d*^ = 0.2, *µ* = 0.7, *σ* = 0.25) respectively. We considered two settings (Supplementary Table S2) by varying the magnitude of the stimulus effect with the weight of the activity GEPs to capture the cases that do (setting 1) and do not (setting 2) require data integration.

## 3 Results

### 3.1 The multilayer graph (MLG) clustering algorithm

We first present a detailed illustration of MLG on *Kowalczyk 1* dataset (hemotapoitic stem and progenitor cells from young and old mice). The first and second rows in Figure 1B depict the two-dimensional visualizations of the low-dimensional embeddings of the *Kowalczyk 1* dataset with t-SNE [33] labelled by condition and cell type, respectively. The third row presents the Louvain clustering labels for each embedding. A direct comparison of the second and third rows illustrates the inaccuracies of each method for cell type identification. To evaluate the performances of these low-dimensional embeddings independent of clustering, we followed a supervised approach. We leveraged a k-nearest neighbor (knn) classifier to classify the cells, and computed the commonly used metric adjusted Rand index (ARI, [34]) between predicted class labels and the true labels in Figure 1C. The ARI values from the knn classification represent the best achievable performances with these low-dimensional embeddings. In addition to this, we also quantified the clustering performance of each method by ARI.

The ARIs of the knn classifier with PCA and cNMF factors are 0.84 and 0.76, whereas the ARI of knn classifier with Seurat and Liger integrated data are 0.75 and 0.70, respectively. While PCA and cNMF keep cells from the same cell type close to each other in their respective low-dimensional spaces, Seurat and Liger perform similarly well in aligning data across conditions. However, all four methods exhibit cell type mix-up and, as a result, none of the methods have an ARI higher than 0.51 when these low-dimensional embeddings are clustered (Figure 1C). In the context of graph clustering, cells are partitioned so that cells within the same cluster are densely connected, while cells in different clusters are loosely connected. We observe for this dataset that, with dimension reduction without data integration, the clustering algorithm tends to separate cells under different conditions, e.g., old and young LT-HSCs with cNMF factors. However, after Seurat and Liger data integration, graph clustering tends to separate MPP cells into different clusters. To leverage strengths of different dimension reduction strategies, MLG clustering first constructs shared nearest neighbour graphs (SNNs) from each of the low-dimensional embeddings. Then, it aggregates the adjacency matrix of the resulting graphs with a union operation and employs modularity maximization [13] to cluster the resulting graph. By aggregating the SNNs obtained from each dimension reduction approach, MLG boosts the signal-to-noise ratio and improves the ARI from individual methods by 25% to 0.76.

In the following sections, we first provide explicit examples supporting the analytical underpinnings of MLG clustering as outlined in the Methods section and demonstrate how different low-dimensional embeddings can complement each other. We then evaluate MLG clustering on both simulated and benchmark datasets and compare it with state-of-the-art methods. We then showcase MLG in two separate scRNA-seq applications involving cells from two [19] and four different conditions [20], respectively. Finally, we showcase how MLG can also be adapted beyond analysis of scRNA-seq to SNARE-seq [21] which profiles transcriptome and chromatin accessibility from the same cells simultaneously.

### 3.2 Aggregating signal from shared nearest neighbour graphs of multiple low-dimensional embeddings boosts the “signal-to-noise” ratio

A challenging aspect of generating low-dimensional embeddings of scRNA-seq data across multiple conditions is that, different dimensionality reduction methods might capture different aspects of the data. As a result, graph representations of the data constructed for downstream clustering might vary significantly between different lower-dimensional embeddings. In the HSCs of old and young mice (*Kowalczyk 1* from Figure 1), SNN graphs constructed from PCA and cNMF factors of the gene expression count matrix (with and without data integration across conditions) have small proportions of overlapping edges (Figure 2A). Specifically, SNN graphs from any two low-dimensional embeddings, except the pair PCA and Seurat-integration, which also performs PCA as the final dimension reduction, have at most 30% of their edges overlapping, indicating low dependence between these constructions. One possible reason for this, as we argue analytically in the Supplementary Materials, is that dimension reduction alters neighbors of cells. Furthermore, we also observe that all of the constructions tend to be sparse as indicated by the proportion of cell pairs with no edges in the individual SNN graphs (bar labelled as “with no edge” in Figure 2B).

**Figure 2.**
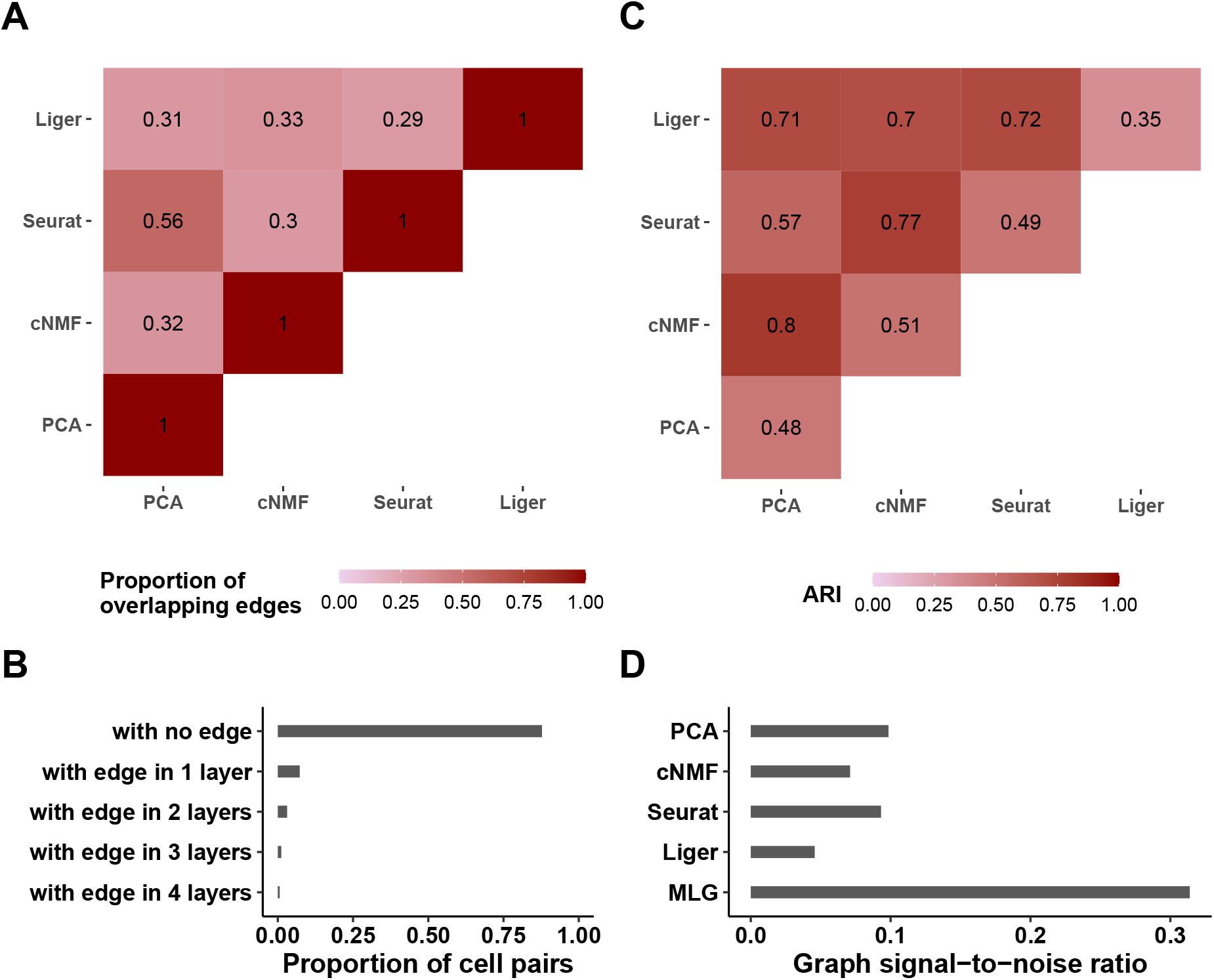
Graph characteristics of SNN graphs and the multilayer graph of the *Kowal-czyk 1* dataset. (A) Heatmap of the proportion of overlapping edges between pairs of SNN graphs from different low-dimensional embeddings. (B) Proportion of cell pairs with edges across aggregation of individual SNN graphs from PCA, cNMF, Seurat and Liger low-dimensional embeddings. The number of layers represents the total number of individual SNN graphs that harbor edges between the cell pairs. (C) ARI of MLG clustering constructed by pairs of SNN graphs from the low-dimensional embeddings indicated as the rows and columns. Diagonal entries represent ARIs of SNN graph clustering from individual low-dimensional embeddings. (D) Estimated graph signal-to-noise ratios of individual SNN graphs and their multilayer graph.

Leveraging these empirical observations and recent advancements in stochastic block models, we show in the Supplementary Materials that aggregating two sufficiently sparse graphs with independent edges by taking union of their edges leads to a graph with amplified “signal-to-noise” ratio. Here, as we presented in the Methods section, we are using a notion of “signal-to-noise” that refers to difference in connectivity of cells that are within the same cluster (i.e., cell type/stage) versus that are in different clusters. Figure 2C presents ARI values of MLG clustering based on pairs of SNN graphs (off-diagonal entries) and those of clusterings based on individual SNN graphs (diagonal entries). We observe that all the 2-layer MLG applications have improved clustering performance compared to SNNs from individual low-dimensional embeddings (e.g., Liger alone achieves an ARI of 0.35 whereas MLG with Liger and PCA low-dimensional embeddings as its two layers achieve 0.71). As expected, since the SNN graphs constructed with PCA and Seurat have more overlapping edges (56%), their 2-layer MLG result in the least improvement in ARI with a value of 0.57 compared to 0.48 and 0.49 with PCA and Seurat alone.

A majority of cell pairs (88%) do not have edges in any of the four layers of the SNN graph constructed from aggregation of PCA, cNMF, Seurat-integration, and Liger SNN graphs (Figure 2B), which is due to sparsity of individual SNN graphs. Furthermore, about 67% percent of the cell pairs with edges, have edges in only one layer, suggesting that different dimension reduction methods are capturing distinct features of the data. Aggregating the four layers with union operation increases the signal-to-noise ratio to three times of any single SNN graph layer (Figure 2D). A direct impact of this is increased clustering accuracy by MLG compared to clustering of SNN graphs from individual low-dimensional embeddings. In fact, MLG clustering almost achieves the accuracy of supervised knn classifiers (0.76 versus 0.80, Figure 1C).

### 3.3 Increased “signal-to-noise-ratio” by MLG aggregation translates into significant improvements in clustering accuracy and stability in computational experiments

To systematically evaluate the ability of MLG in improving clustering performance over clustering with individual graphs from specific low-dimensional embeddings, we conducted simulations. Specifically, we considered the general simulation setting from [22] where cell types exhibited condition specific effects and devised two settings by varying the condition effect size. We generated 100 simulation replicates for each setting, where each replicate included a total of 2,500 cells from 3 cell types and across 2 conditions. We adapted the simulation pipeline from Splatter [32] and re-implemented it based on the Python code from [22] to account for differential expression between conditions.

We considered four different dimension reduction procedures for MLG: PCA and cNMF, which do not perform comprehensive data integration, Seurat and Liger both of which perform data integration for cells from different conditions. We varied the apparent parameters of each method such as the numbers of PCA and NMF components and numbers of neighbours in the construction of SNN graphs. To compare with the MLG clustering, we applied both k-means and Louvain clustering on the SNN graphs constructed by the individual low-dimensional embeddings from these methods. Furthermore, we employed a supervised k-nearest neighbor classifier to establish the best achievable clustering performance for each graph in terms of ARI.

We first assessed the level of dependence between the SNN graphs constructed from these four low-dimensional embeddings across the simulation replicates and observed that majority of the cell pairs were connected only in one of the SNN graphs, resulting in only 3.6% of the cell pairs common to all 4-layers, a level comparable to those of the benchmark datasets (Figure 3A). Furthermore, the estimated signal-to-noise ratios of each graph supported the signal boost by MLG (Figure 3B). Figure 3D summarizes the ARI values of supervised knn, Louivan and k-means clustering with each individual low-dimensional embedding as a function of the numbers of PCA/cNMF components across the simulation replicates. We observe that MLG provides a median increase of 13%, 9%, 34%, and 55% in ARI compared to Louvain clustering of individual PCA, cNMF, Seurat, and Liger-based low-dimensional embeddings, respectively (first row of Figure 3D). Improvement in ARI by the MLG clustering with the Louivan algorithm is even higher (median levels of 85%, 9%, 80%, and 45%) compared to k-means clustering of the low-dimensional embeddings by each of the four methods (first versus second rows of Figure 3C). Furthermore, MLG yields accuracy levels that are comparable to those of best knn accuracy in a supervised setting by these four methods (third row of Figure 3D). Since both the dimension reduction methods and SNN graph construction depend on key parameters such as the numbers of latent factors and numbers of neighbours, we varied these parameters in a wide range and observed robustness of MLG to the choice of these two parameters (Figure 3C, D).

**Figure 3.**
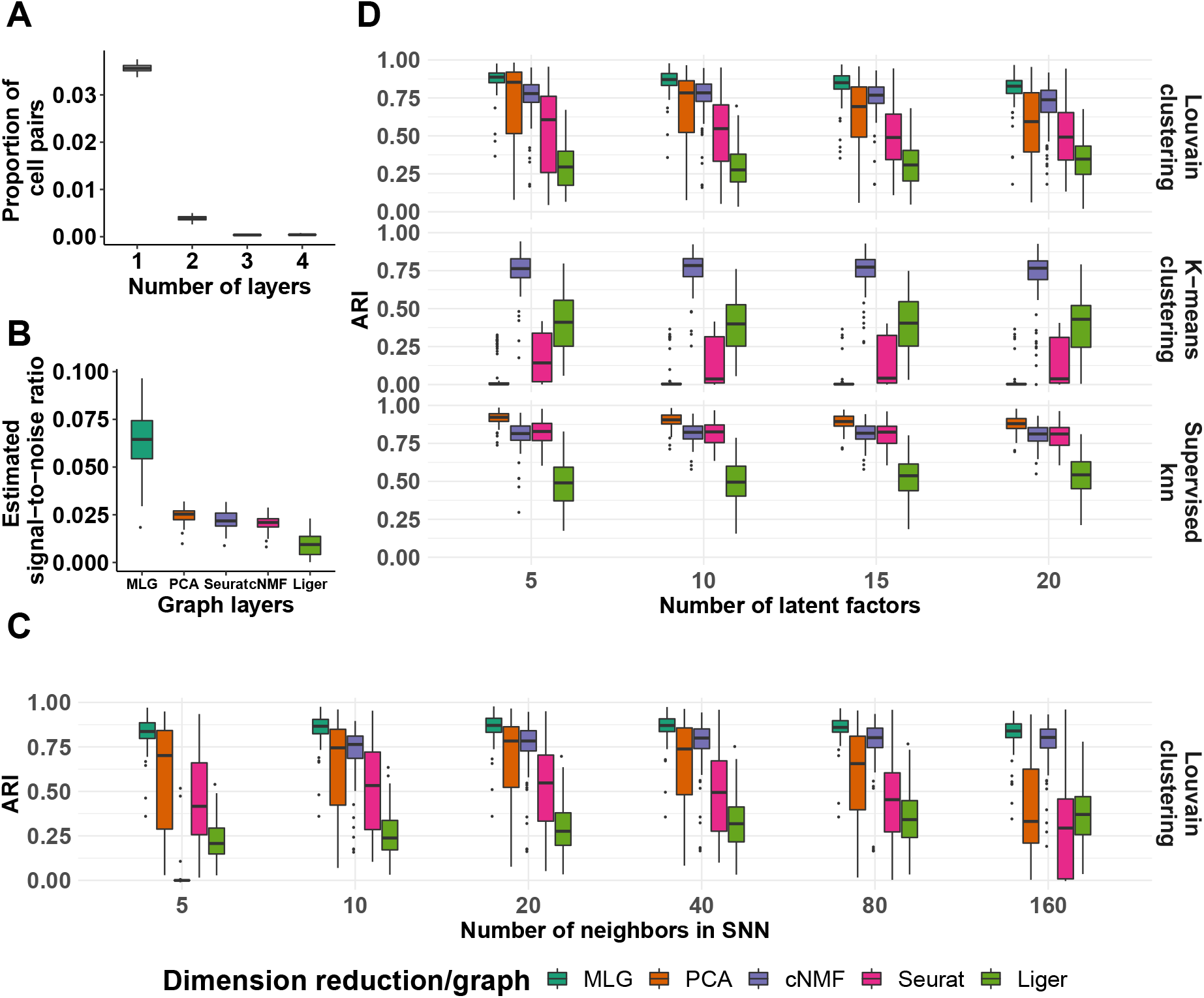
Simulation results for the “large-condition-effect” setting. (A) Proportions of cell pairs with edges across different numbers of layers of MLG constructed from SNN graphs of PCA and cNMF, Seurat, and Liger (with 20 neighbours in SNN graphs and 10 latent factors in low-dimensional embeddings). The boxplots depict the proportions across all the simulation replicates. (B) Estimated signal-to-noise ratios of SNN graphs constructed from different low-dimensional embeddings and their multilayer graph across all the simulation replicates (with 20 neighbours in SNNs and 10 latent factors in low-dimensional embeddings). (C) Louvain clustering accuracy of SNN graphs as a function of numbers of neighbors in SNN graph construction (with 10 latent factors in low-dimensional embeddings). (D) Adjusted Rand index comparison of Louvain and k-means clustering of SNN graphs from different low-dimensional embeddings and their MLG as a function of number of latent factors in the low-dimensional projections (with 20 neighbours in SNN graphs). ARI values of supervised knn classifiers for individual SNN graphs are provided as reference.

### 3.4 MLG clustering outperforms other methods in recovering known biological signal

We next compared MLG clustering to state-of-the-art scRNA-seq dimension reduction and data integration methods on the benchmark datasets with cells from multiple conditions and ground truth cell type labels, namely *Kowalczyk 1, Kowalczyk 2, Mann*, and *Cellbench* (Figure 4). These datasets exhibit different levels of difficulty for clustering based on the average mixing metric [7], which ranges from 5 to 300 and quantifies how mixed different group of cells are (Figure 4A). The separations between conditions are relatively small in datasets *Kowalczyk 1, Kowalczyk 2, Cellbench*, indicating that condition specific responses of cells do not dominate over their cell type-specific expression programs, and fairly large in *Mann* with average mixing metric of 10.3, 12.0, 13.7, and 27.0, respectively. In contrast, separations between cell types are low in *Mann* and high in *Cellbench*, with average mixing metric of 41.8 and 278.7, respectively. This exposition of condition and cell type separation levels indicate that *Cellbench* is the easiest to cluster and *Mann* is the hardest. In these benchmarking experiments, we extended the set of methods we considered to include Harmony integration, scVI batch correction, and scAlign integration in addition to PCA and cNMF dimension reduction, Seurat integration, Liger integration. We performed both k-means clustering with the latent factors estimated from these dimension reduction/data integration methods and also Louvain clustering with graphs constructed from their low-dimensional embeddings (with package default weighted SNN for Seurat, package default SFN graph for Liger, unweighted SNN graphs for all other methods). As an overall measurement of the difficulty of the clustering task, we performed supervised knn classification using SNN/SFN graphs and observed largely similar supervised knn accuracy from different low-dimensional embeddings (Figure 4B) (with scAlign on *Cellbench* and *Mann* datasets as notable exceptions). The accuracies of k-means and Louvain clustering with graphs from individual low-dimensional embeddings are markedly lower than their corresponding knn classifiers for datasets *Kowalczyk 1, Kowalczyk 2*, and *Mann*. Harmony, Seurat and PCA perform well for *Cellbench* which is an easier dataset in terms of clustering since the separation between the cell types is large (Figure 4A). MLG clustering outperforms the alternatives across all datasets with a minimum of 13% and a maximum of 64% increase in ARI. Furthermore, as apparent from the Louvain clustering results of individual low-dimensional embeddings in Figure 4B, MLG does not require each individual SNN graph it aggregates over to perform well. For example, while Seurat, Liger, PCA and cNMF have individual ARIs of 0.35, 0.11, 0.10, 0.00, respectively, by combining SNN graphs resulting from these, and boosting signal-noise-ratio, MLG outperforms their individual performances with an ARI of 0.51 for the *Mann* dataset that appears to be the most challenging to cluster.

**Figure 4.**
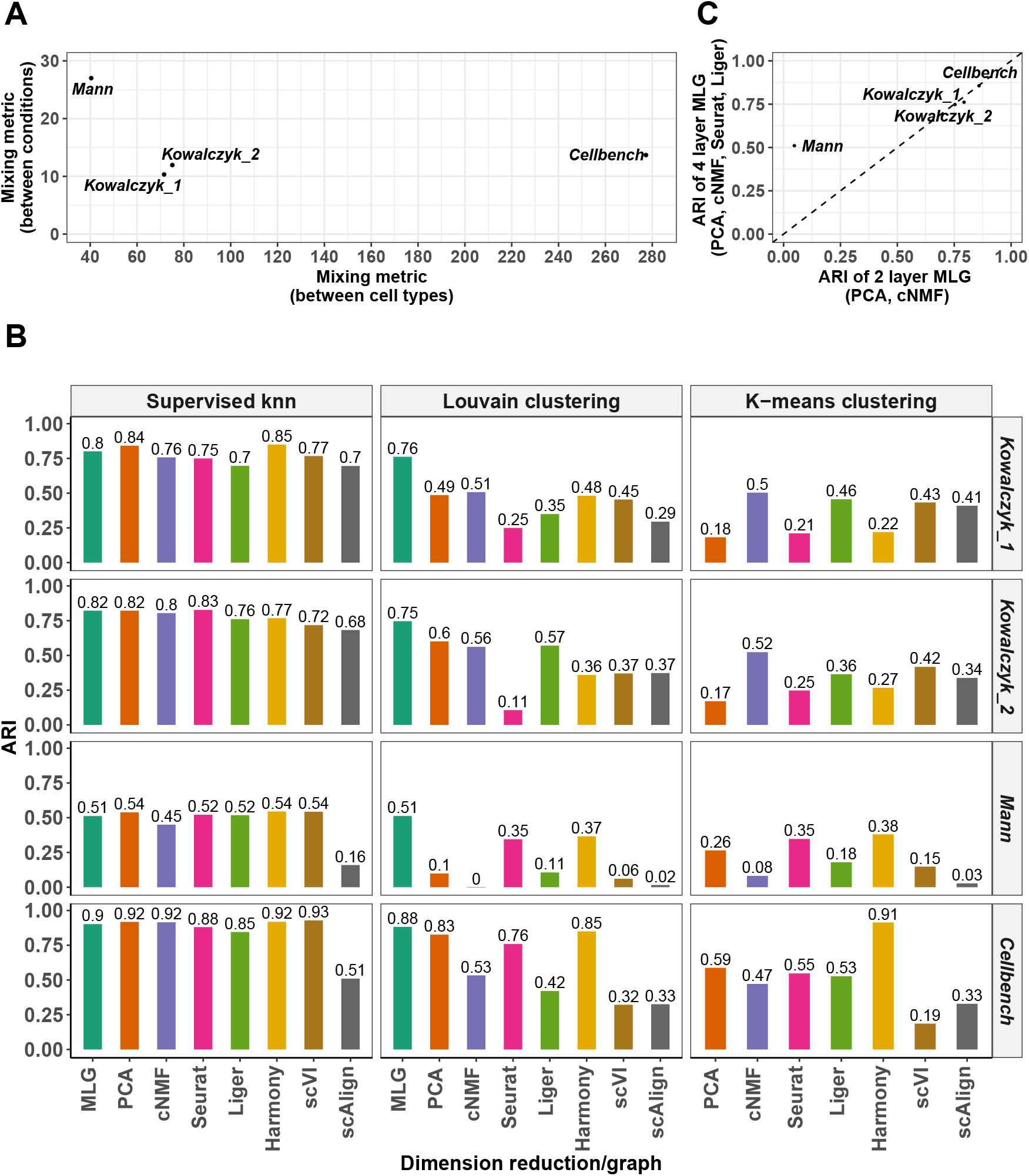
Evaluation with benchmark datasets. (A) Average mixing metric between cell types (x-axis) and conditions (y-axis) for the four benchmark datasets. The mixing metric ranges between 5 to 300, with larger values indicating large separation between the groups (e.g., cell types or conditions). (B) Louvain and k-means clustering accuracy of different methods across the benchmark datasets. Supervised knn classification for SNN graphs from each low-dimensional embedding is provided as a measure of the difficulty of the clustering task. (C) Comparison of the 4-layer MLG (PCA, cNMF, Seurat-integration, Liger) with the 2-layer MLG (PCA, cNMF) across the benchmark datasets.

We leveraged these benchmark datasets to further investigate the impact of different low-dimensional methods aggregated by MLG. Figure 4C displays the ARI values of 2-layer MLG which aggregates over only PCA and cNMF and 4-layer MLG which also aggregates over low-dimensional embeddings from data integration with Seurat and Liger. Empirically, when condition separation, as measured by the average mixing metric and also visualized by tSNE or UMAP visualizations of the data prior to integration, is low as in *Kowalczyk 1, Kowalczyk 2*, and *Cellbench* datasets, the two MLG strategies perform similarly; however, for datasets with larger separation between conditions (*Mann*), MLG benefits significantly from aggregating over low-dimensional embeddings from data integration.

### 3.5 MLG strategy is robust to imbalanced cell type representations across conditions

When considering scRNA-seq datasets across multiple conditions, a key challenge is the ability to identify distinct cell types in cases with varying levels of representation of cell types under different conditions. Among the benchmark datasets we considered, *Kowalczyk 1* and *Mann* have relatively balanced cell types in different conditions. Specifically, in the *Kowalczyk 1* dataset, the three different cell types (LT-HSC, MPP, ST-HSC) vary at proportions of (0.17, 0.18, 0.18) and (0.16, 0.16, 0.15) among the old and young mice, respectively. Similarly, in the *Mann* dataset, the three different cell types (LT-HSC, MPP, ST-HSC) vary at proportions of (0.11, 0.02, 0.09), (0.12, 0.05, 0.12), (0.04, 0.03, 0.07), and (0.10, 0.10, 0.14) among the old-no-stimuli, old-stimuli, young-no-stimuli, and young-stimuli mice conditions, respectively. Taking advantage of these balanced datasets, we conducted a computational experiment to investigate performance under imbalanced cell type representations across the conditions. Specifically, we subsampled “old MPP”cells in *Kowalczyk 1*, and “old-stimuli-MPP” cells in *Mann*, at proportions 0, 0.10, 0.25, 0.50, 0.75 and 1 of the original size, where 1 corresponded to the original dataset. Figure 5A reports the mean, 5, and 95 percentiles of the ARI values across 20 sub-sampling replications for the 2-layer MLG (PCA and cNMF), 4-layer MLG (PCA, cNMF, Seurat, Liger), and Louvain and k-means clustering with PCA, cNMF, Seurat, Liger, Harmony latent factors. For *Kowalczyk 1*, 2-layer MLG outperforms other methods and is robust to the imbalance of the cell types between conditions. PCA and cNMF can accommodate the small separation between conditions of this dataset without any explicit data integration (Figure 4B). However, methods utilizing data integration are affected by the misalignment of cells, which in turn reduces their clustering accuracy (Seurat, Liger). The 4-layer MLG is also relatively robust to cell type imbalance despite its aggregation over Seurat and Liger, and outperforms clustering with all individual low-dimensional embeddings in accuracy and stability. Since the *Mann* dataset has larger separation between conditions, aggregation over only methods without data integration (2-layer MLG) results in similarly poor performance as PCA and cNMF. In contrast, the 4-layer MLG, by aggregating over data integration methods Seurat and Liger in addition to PCA and cNMF, has the highest ARI. Compared to the balanced case (labeled as 1 in Figure 5A), 4-layer MLG suffers from accuracy loss because of mis-aligned cells in integration; however, it still performs better than using just one layer of integrated data.

**Figure 5.**
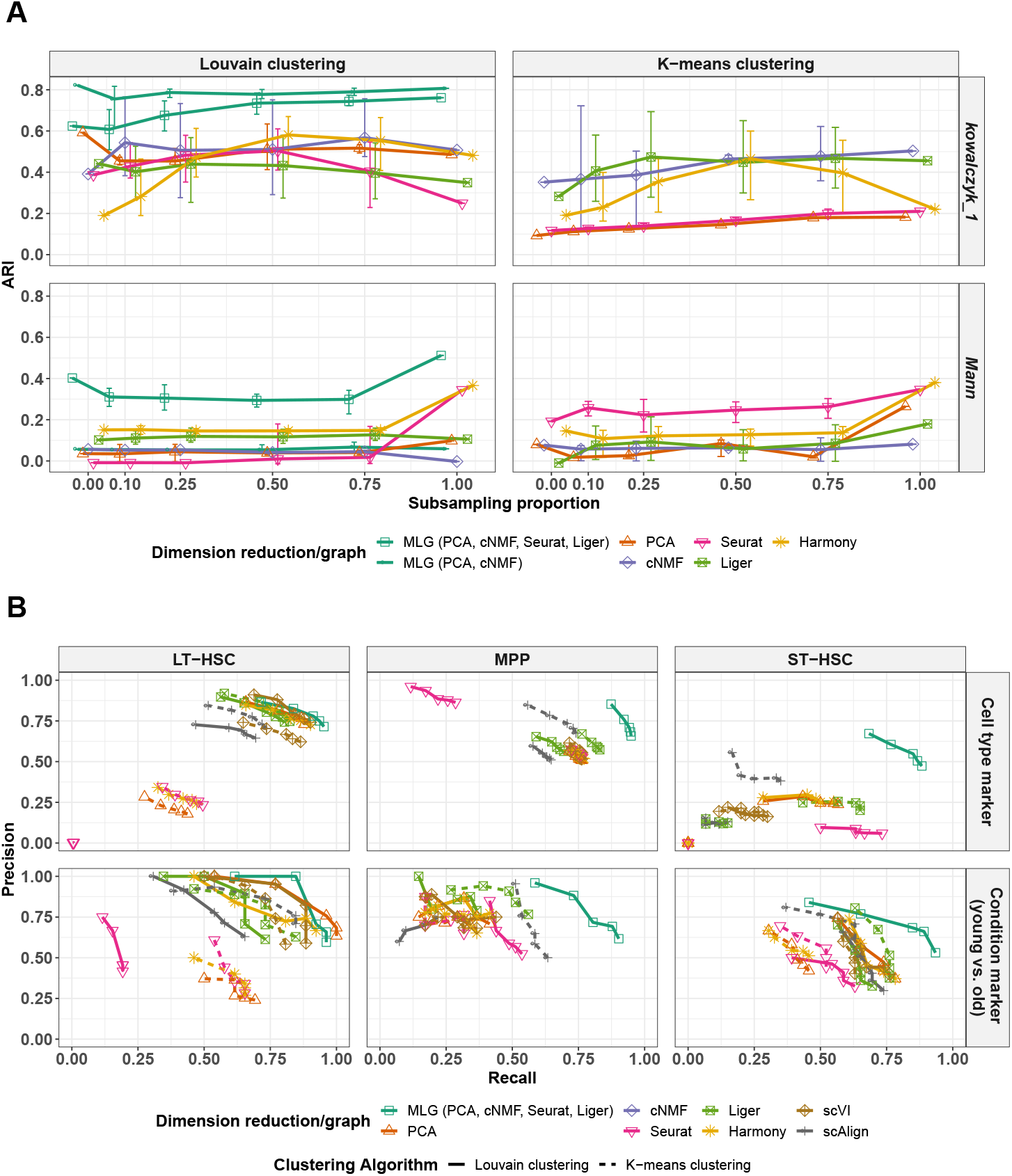
Robustness analysis of clustering over imbalanced samples and downstream impact of MLG clustering on marker gene identification. Adjusted Rand index from Louvain and k-means clustering of low-dimensional embeddings with varying levels of imbalance in the proportion of cells from different cell types. For the *Kowalczyk 1* and *Mann* datasets, varying proportions of MPP cells from the “old” and “old, stimulated” conditions were subsampled for the respective analysis. Reported ARI values correspond to mean, 5 and 95 percentiles across 20 subsampling experiments. (B) Differential expression analysis for cell type/cluster marker gene identification and condition (young vs. old) DE gene identification in the *Kowalczyk 1* dataset. The precision-recall (PR) values in the top panel evaluate inferred cluster (i.e., cell type) marker genes of each method against cell type marker genes defined with ground truth cell type labels as gold standard. The bottom panel evaluates the inferred condition (i.e.,age) DE genes of each cell type against the gold standard. Gold standard cell type marker and age DE genes are defined as genes with Bonferroni corrected p-values less than 0.01 in the differential expression analysis with ground truth cell labels. The PR values are reported at cutoffs of 0.2, 0.1, 0.05, 0.01, and 0.001 for Bonferroni adjusted p-values in the cluster marker gene and age DE gene identification analysis.

### 3.6 MLG clustering across multiple stimuli leads to more powerful downstream differential expression analysis

We evaluated the impact of improvement in clustering accuracy by MLG clustering on the down-stream analysis of identifying marker genes of individual cell types and cell type-specific differentially expressed (DE) genes across conditions. We first generated “gold standard” marker genes and lists of differentially expressed genes using the true labels of the cells in the benchmark datasets (Methods). Next, we identified cluster-specific marker genes and lists of DE genes across conditions using the cluster assignments obtained with MLG clustering and the alternative methods used for the benchmark datasets. We evaluated whether more accurate separation of the cell types by MLG leads to marker and DE gene identification that aligns better with the gold standard sets with precision-recall (PR) curves (Figure 5B for the *Kowalczyk 1* datasets, Supplementary Figures S2, S3 for the *Kowalczyk 2* and *Mann* datasets). Overall, MLG exhibits better PR values for both cell type marker and condition DE gene identification across multiple datasets. More specifically, MLG is the only method with moderate to high precision-recall in identifying cell type marker genes for ST-HSC, whereas all other methods perform poorly in identifying marker genes specific to this cell type (Figure 5B).

### 3.7 MLG clustering confirms disrupted differentiation patterns in HSPCs lacking the murine *Gata2* −77 enhancer

We utilized MLG clustering to analyze scRNA-seq data of hematopoietic progenitor cells sorted from E14.5 fetal livers of *−*77^+*/*+^ (wild type, WT) and *−*77^*−/−*^ (mutant, MT) murine embryos (dataset *Johnson20* from [19]). The mutant condition corresponded to homozygous deletion of the murine *Gata2* −77 enhancer, located −77kb upstream of the *Gata2* transcription start site, as described in [19]. The samples from both WT and MT mice included a complex mixture of progenitors with diverse transcriptional profiles [35] from a pool of common myeloid progenitor (CMP) and granulocyte-monocyte progenitor (GMP) cells and resulted in 14, 370 cells after pre-processing [19]. Exploratory analysis with the data visualization tools t-SNE [33], UMAP [36], and SPRING [37] revealed a small separation between the cells from the wild type and mutant conditions and an average mixing metric of 15.25. Furthermore, Johnson et al. [38] previously showed that *−*77^+*/*+^ fetal liver CMPs have the potential to differentiate into erythroid and myeloid cells ex vivo. In contrast, the mutant *−*77^*−/−*^ CMPs and GMPs generate predominantly macrophage progeny. In this context, the differential heterogeneity of the wild type and mutant progenitor populations necessitates single-cell transcriptional analysis to establish mechanistic insights. This instigated us to proceed with clustering of the cells without data integration since the subsampling experiments with benchmark datasets highlighted that data integration may cause misalignment of cells in this setting with potentially imbalanced cell types.

We constructed a multilayer graph from PCA (after regressing out total counts, mouse batch effects, and percentage count of mitochondrial transcripts) and cNMF factors. Figure 6A displays the SPRING visualization of the MLG clustering with four optimally chosen clusters based on the eigen-gap heuristic [23], and also highlights pseudo-time trajectories of the wild type and mutant cells. We linked these clusters to established cell populations as (1) CMP, (2) erythroid/megakaryocyte, (3) bipotential GMP and monocyte, (4) neutrophils, by leveraging established lineage defining markers of these HSPC populations [39] i.e., *Flt3* and *Hlf* for CMPs, *Hba-a2* and *Car1* for erythrocytes; *Ly86* and *Csf1r* for monocytes; *Gstm1* and *Fcnb* for neutrophils, and *Pf4* for megakaryocytes (Figure 6A). Overall, cells in different clusters exhibited expression patterns of documented lineage markers [39] consistent with their inferred cluster labels (Supplementary Figure S6). Furthermore, an unbiased marker gene analysis of each MLG cluster with MAST [40] (Supplementary Figure S5) yielded marker genes. A gene set enrichment analysis with top 50 marker genes of each cluster agreed with the MLG cluster labels based on known lineage markers (Supplementary Figures S11).

Next, we compared MLG clusters with Louvain clustering of the cell graphs constructed from PCA, cNMF factors, Seurat-integration, and Liger (Figure 6B). cNMF, Seurat-integration, and Liger tended to partition the cells based on their mitochondrial gene expression. Specifically, the clusters in cNMF, Seurat-integration and Liger are driven by the mitochondrial gene expression patterns apparent in the SPRING plot (Supplementary Figure S4). Concordant with these partitionings, the top 10 marker genes for cNMF cluster 1, Seurat cluster 4, and Liger cluster 2 have 5, 7, and 7 mitochondrial genes, respectively. While it is possible to adjust for mitocondrial gene expression of the cells for some settings such as PCA, cNMF and Liger do not encompass mechanisms to adjust for potential continuous confounders and the Seurat-integration framework does not enable adjustment for biological variables in its CCA factors. While the clustering of the low-dimensional embedding from PCA is similar to the MLG clustering, it merges cells expressing *Gata2* and erythroid specific-gene *Car1* with cluster 1 that represents early progenitors (Supplementary Figures S6, S7) and results in a markedly smaller cluster of erythroid cells. In addition to these differences, established lineage markers do not delineate clusters from cNMF, Seurat-integration and Liger as expressing specific cell type specific markers as clearly as MLG clustering (Supplementary Figures S7, S8, S9, S10). Collectively, this analysis illustrates the power of MLG in identifying cell types/stages with low signal by its aggregation strategy.

**Figure 6.**
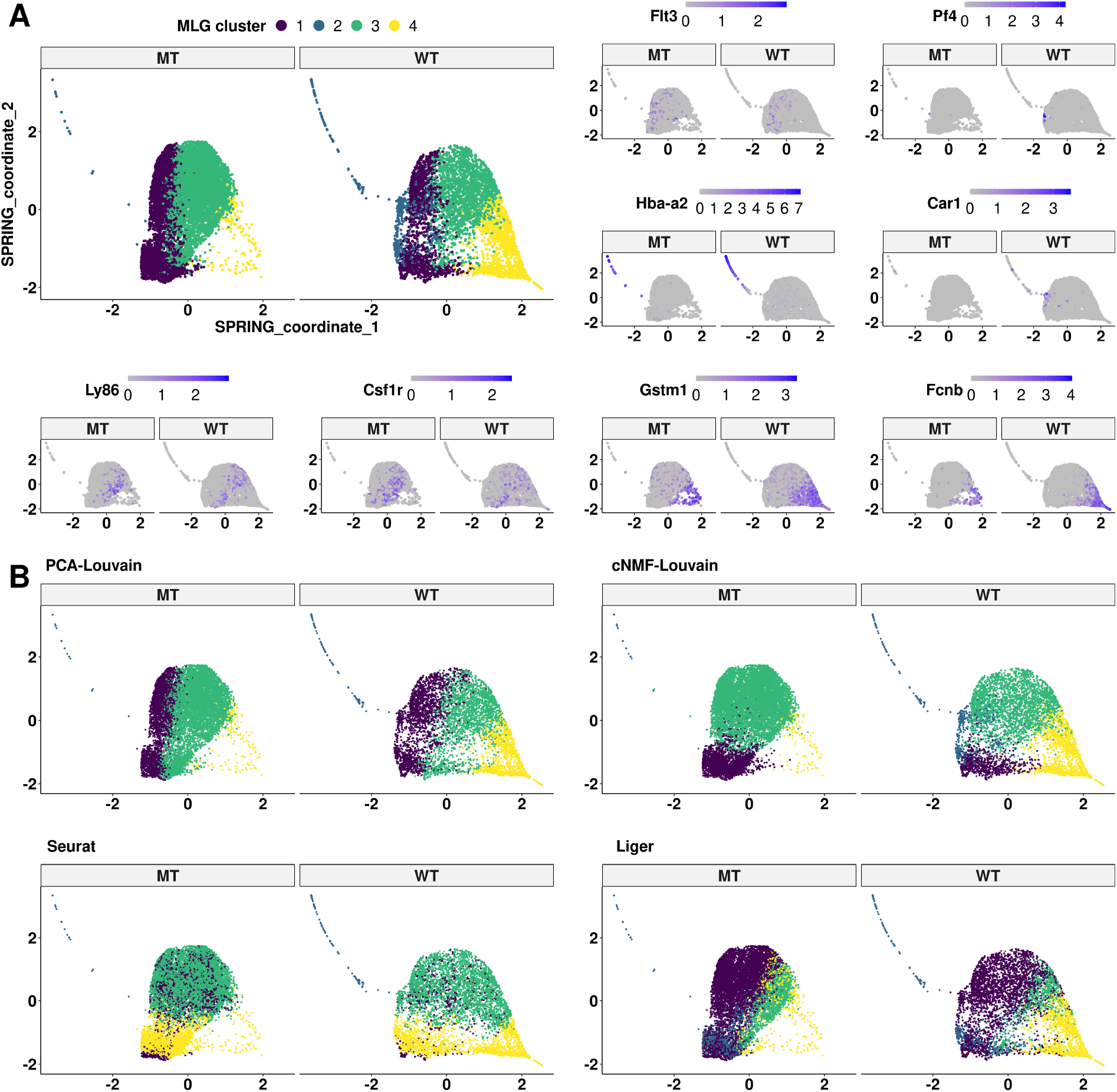
Clustering analysis of the *Johnson20* [19] dataset. (A) SPRING [37] visualization of mutant and wild type cells from *Johnson20*. Cells are labeled with MLG clustering (top left) and expression of linage marker genes. (B) SPRING visualization of mutant and wild type cells with cell labels from Louvain clustering of SNN graphs with PCA, cNMF factors, Seurat integrtion, and Liger low-dimensional embeddings.

### 3.8 MLG uncovers cell stages in mouse HSPC under four experimental conditions

Dataset *Muench20* [20] contains scRNA-seq of 813 mouse hematopoietic progenitor cells under 4 conditions: wild type (*Gif* 1^+*/*+^), heterozyous *Gfi1* R412X mutation (*Gif* 1^*R*412*X/−*^), homozygous *Gfi1* R412X mutation (*Gif* 1^*R*412*X/R*412*X*^), heterozygous *Gfi1* R412X mutation and one silenced *Irf8* allele (*Gif* 1^*R*412*X/−*^*Irf* 8^+*/−*^). Joint dimension reduction was conducted on all the cells, with PCA, cNMF, Seurat integration, and Liger, followed by SNN graph construction with each individual dimension reduction results. Louvain graph clustering was applied to each SNN graph along with MLG with these SNN graphs as four layers. A total of seven clusters were identified with the eigengap heuristic [23] (Figure 7B). Through the expression patterns of HSPC-relevant transcription factors and granule protein-encoding genes (Figure 7A gene set 1), surface marker genes (Figure 7A gene set 2) and linage marker genes (Figure 7A gene set 3), we labelled the clusters 1 to 7 as HSPC, Multi-Lin (multi-lineage progenitor), Mono-1, Mono-2 (Monocyte progenitor), Neu-1, Neu-2, Neu-3 (Neutrophil progenitor). The MLG clustering results also matched the author annotated cell labels from [20] with the exception of the extremely small clusters, e.g. NK(3 cells), DC(5 cells), missed by MLG. We compared the resulting clustering from each method using the author annotated cell labels as gold standard. Since there are 17 distinct cell labels in the author annotation set, we used a wide range of numbers of clusters in the comparison (Figure 7C). MLG yielded clustering results most closely in agreement with the gold standard cell labels. Moreover, it performed more robustly against the choice of numbers of clusters, highlighting the overall robustness of the MLG aggregation strategy.

**Figure 7.**
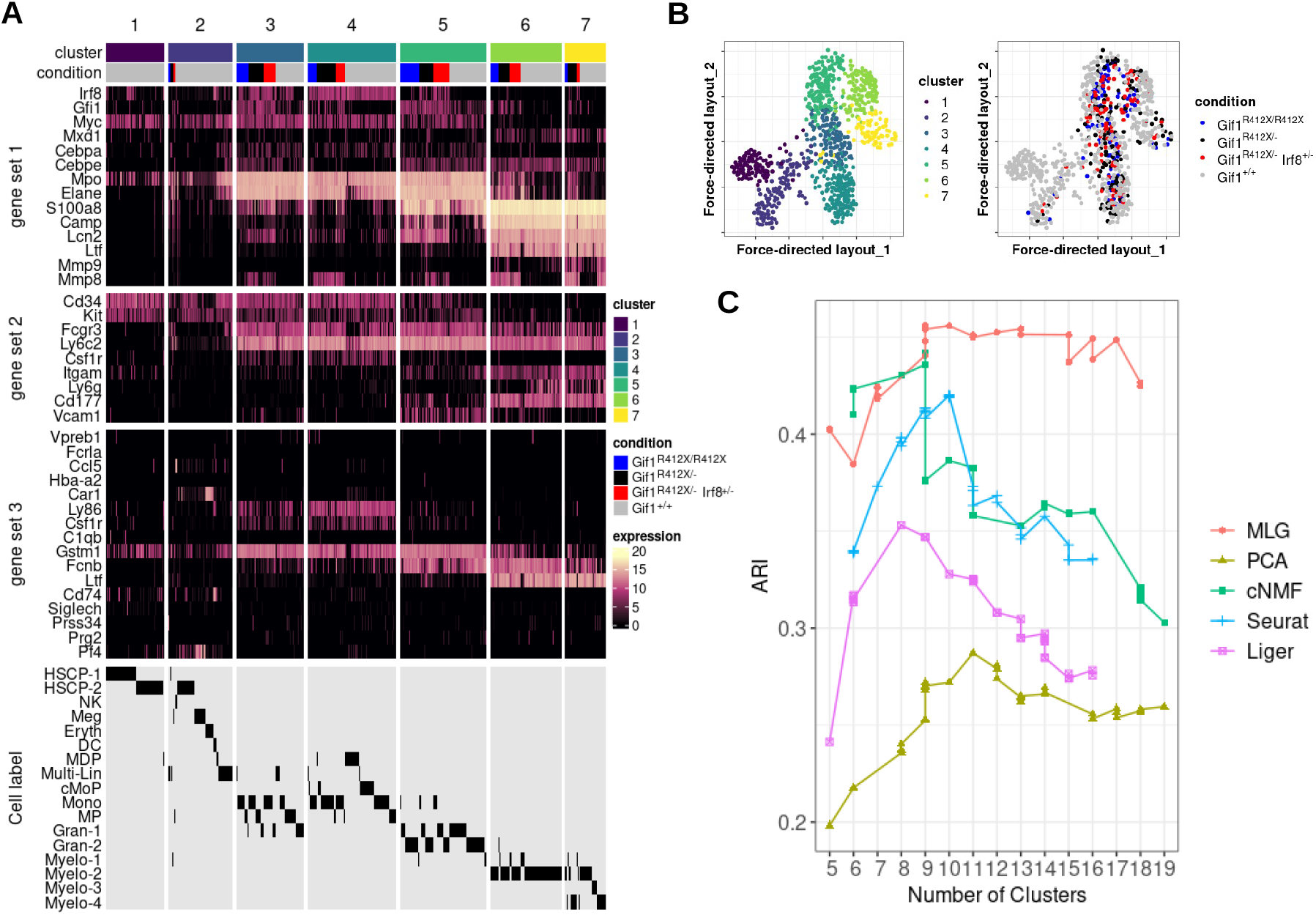
Clustering analysis of the *Muench 20* [20] dataset. (A) Heatmaps of the cell-level gene expression for HSPC-relevant transcription factors and granule protein-encoding genes (gene set 1), surface markers (gene set 2) and lineage defining markers (gene set 3). Cells are grouped by MLG clustering and cell conditions. MLG clustering is compared with author annotated cell labels from [20] in the last row of the heatmap. (B) Visualization of the dataset with the force directed layout of MLG. Cells are labeled with MLG clusters (left) and cell conditions (right). (C) ARIs are computed between author annotated cell labels and clustering results of MLG and individual SNN graphs constructed from low-dimensional embeddings by PCA, cNMF, Seurat-integration and Liger.

### 3.9 MLG provides stable and separable clusters for the analysis of SNARE-seq

Next, we showcase broader applicability of MLG to multi-modal sequencing data with a SNARE-seq dataset that profiles transcriptome (snRNA-seq) and chromatin accessibility (snATAC-seq) in the same cell. The *Chen19* dataset [21] contains SNARE-seq profiles of cells from neonatal and adult mouse cerebral cortices. We used Seurat and Liger for dimension reduction of snRNA-seq and utilized Latent Semantic Indexing [30] (LSI) for dimension reduction of snATAC-seq. Each pair of low-dimensional embeddings of snRNA-seq and snATAC-seq were combined to generate a weighted nearest neighbor (WNN) graph [31]. These resulting graphs were then inputted to MLG (Figure 8A).

**Figure 8.**
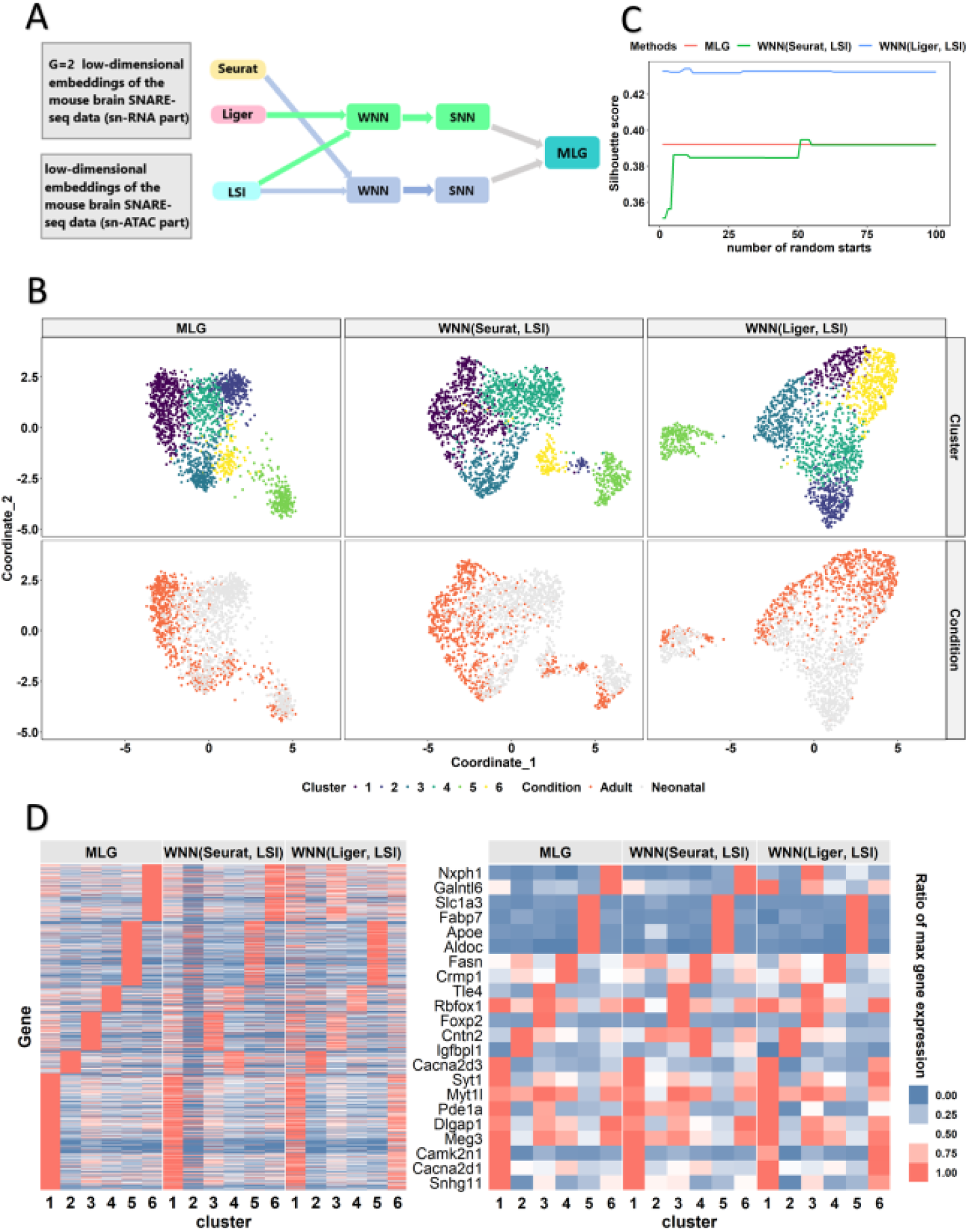
Application to SNARE-Seq [21] dataset *Chen19*. (A) Data processing workflow of MLG for SNARE-seq. Dimension reduction methods Liger and Seurat are applied to snRNA-seq, Latent Semantic Indexing (LSI) [30] is applied to snATAC-Seq. Weighted nearest neighbor (WNN) graphs [31] are constructed using Liger-LSI and Seurat-LSI low-dimensional embeddings. Finally, the two WNN graphs are inputted to the MLG framework. (B) Visualization of clusters and conditions on the force-directed layout of MLG and the UMAP coordinates of WNN(Seurat, LSI) and WNN(Liger, LSI). (C) Average silhouette scores for the three clustering methods, computed with the corresponding UMAP or force-directed layout coordinates. X-axis is the number of random starts inputted to the Louvain algorithm. (D) Heatmap of the scaled gene expression for all the genes (left) and selected genes (on the right). Analysis of another set of replicates (replicate 2) from the *Chen19* dataset yielded similar results as the replicate 1 presented here (Supplementary Materials Figure S12).

We compared the MLG clustering results with those of individual WNN graph clustering in terms of separability, as measured by the average silhouette score, and algorithmic stability, as measured by the variation of silhouette score with different starting values in the Louvain algorithm. Taking into account the major cell types, i.e., neurons, endothelial cells, ependymal cells, oligodendrocytes, microglia and astrocytes, in the mouse brain, we expected to detect at least six clusters when maximizing the average silhouette score. While the separability of WNN(Liger, LSI) is slightly higher than those of WNN(Seurat, Liger) and MLG, MLG yields more stable clusters, with a silhouette score invariable to the varying numbers of starting values (Figure8C).

Next, we sought to align MLG clusters to known cell types by leveraging the expression of cell type marker genes. Cells in cluster 1, have high expression in *Meg3* [21], *Snhg11* [41] and *Syt1* [42], which are marker genes for neurons. Cluster 2 is identified to be neuroblast by the expression of *Igfbpl1* [43]. Cells in cluster 5 show an expression pattern consistent with astrocytes based on marker genes *Aldoc* and *Slc1a3* [42]. The other three clusters are formed by subtypes of neurons: cluster 3 and 4 are enriched for *Tle4* and *Crmp1* expression, respectively, suggesting different kinds of excitatory neurons [21]; cluster 6 expresses *Nxph1*, representing inhibitory neurons [44]. In the marker gene heatmap (Figure 8D), MLG and WNN(Seurat, LSI) clusters exhibit more distinguished cluster specific gene expression compared to WNN(Liger, LSI), for which clusters 1 and 6 both have high expression in neuron marker genes.

## 4 Discussion

We have introduced MLG clustering as a versatile method to identify cell types/stages in scRNA-seq data from multiple stimuli by aggregating cell graphs of multiple low-dimensional embeddings of the data. MLG construction capitalizes on the complementary information captured with SNN graphs from different dimension reduction methods such as PCA and cNMF in addition to data integration methods such as Seurat and Liger for scRNA-seq from multiple stimuli. By aggregating sparse SNNs with low edge overlap, MLG amplifes the signal-to-noise ratio. Here the signal-to-noise ratio is quantified by comparing the within cluster connectivity of cells to the between cluster connectivity.

While the MLG clustering framework is flexible in the types of dimension reduction methods it can aggregate over, computational experiments with both simulated and benchmark datasets demonstrated that combining SNN graphs from low-dimensional embeddings of PCA and cNMF perform well when the stimuli is not the dominant effect and it benefits from aggregating over Seurat-integration and Liger when data integration is essential to accommodate the stimuli effect. Furthermore, both PCA and cNMF are fast, robust, and easy to tune compared to other dimension reduction methods for scRNA-seq. Another guiding principle for building MLG is to avoid the simultaneous use of methods that are highly related to prevent false cell-to-cell connections from amplification. The mixing metric performed as a robust quantity for capturing the level of the stimuli effect to guide the aggregation task in terms of whether or not to include low-dimensional embeddings from data integration. Overall, we found that Louvain clustering of MLG exhibited superior performance over clustering with individual low-dimensional embeddings both in accuracy of the clustering and the downstream marker gene identification.

While our focus in this work has been predominantly scRNA-seq datasets from multiple conditions, we showcased how MLG can be applied to analyze multi-modal SNARE-seq data from joint profiling of transcriptome and chromatin accessibility [21]. We expect MLG clustering to be generally applicable for SNARE-seq, Share-seq [45] and other single cell data modalities such as scHi-C [46], joint profiling of transcriptome and DNA methylation [47]. In these applications, each modality yields individual low-dimensional embeddings of the cells and aggregation of their graph products can increase the overall signal.

## 5 Data availability

Datasets *Kowalczyk 1* and *Kowalczyk 2* are available at NCBI GEO with accession number GSE59114. The *Mann* dataset is available at NCBI GEO with accession number GSE100426. The *Cellbench* dataset is available from Github at https://github.com/LuyiTian/sc_mixology/blob/master/data/mRNAmix_qc.RData. The *−*77^+*/*+^ and *−*77^*−/−*^ datasets are available at NCBI GEO with accession number GSE134439. Dataset *Muench20* is available at https://www.synapse.org/#!Synapse:syn16806696. The SNARE-seq dataset, *Chen19*, is available at the Gene Expression Omnibus database under accession number GSE126074.

We implemented the MLG clustering in an open-source R package available on Github https://github.com/keleslab/mlg.

## 6 Acknowledgements

