## Supplementary materials for "MLG: Multilayer graph clustering for multi-condition scRNA-seq data"

#### Aggregation of multiple SNNs boost signal-to-noise ratio

To investigate the impact of aggregating SNNs from multiple low-dimensional embeddings, we first define "signal-to-noise ratio" of a graph. We base our analysis on stochastic block models (SBMs) [1] which are generative random graph models that serve as canonical models for investigating graph clustering and community detection. In the view of SBMs, cells are vertices of a graph and clusters depicting cell types represent distinct communities. We denote the total number of cells (vertices) by $n$, the number of communities (clusters) by $K$. Let $\sigma$ be a cluster assignment function where $\sigma(i)$ corresponds to the cluster label for cell $i$, and $\theta_{\sigma(i)\sigma(j)}$ denote the connectivity probability (i.e., edge probability) between cells $i$ and $j$. Furthermore, let $\theta_{in}$ and $\theta_{out}$ denote the minimum in-cluster and maximum out-of-cluster connectivity probability. We denote the overall parameter space as

$$
\begin{aligned}
\Theta(n, K, \theta_{in}, \theta_{out}, \beta) = \Big\{ \big(\sigma, \{\theta_{k\ell}\}_{k,\ell=1,\dots,K}\big) &: \sigma \in [K]^n, \\
& n_k \in \Big[\frac{n}{\beta K}, \frac{\beta n}{K}\Big], \forall k \in [K], \\
& \{\theta_{kl}\} \in [0,1]^{K\times K}, \text{ and} \\
& \theta_{kk} \geq \theta_{in}, \\
& \theta_{k\ell} \leq \theta_{out} \text{ if } k \neq \ell \Big\},
\end{aligned} \tag{1}
$$

where $n_k$ denotes the size of cluster $k$, $k = 1, \cdots, K$ and $\beta \geq 1$ is a constant setting a lower bound for minimum cluster size.

For a cluster assignment $\hat{\sigma}$, the mismatch rate, which quantifies the discrepancy between the true cluster labels and the assignments from $\hat{\sigma}$, is defined in a way that accounts for equivalent partitions

$$
r(\sigma, \hat{\sigma}) := 1 - \max_{\pi} \frac{1}{n} \sum_{i=1}^{n} \mathbb{1}\{\sigma(i) = \pi(\hat{\sigma})(i)\}, \tag{2}
$$

where $\mathbb{1}$ is an indicator function, $\pi$ is a permutation of cluster labels of $n$ vertices, and the function $\mathbb{1}\{\sigma(i) = \pi(\hat{\sigma})(i)\}$ returns 1 if the cluster assignment for cell $i$ is correct under a clustering (i.e., partition) $\pi(\hat{\sigma})$ which is equivalent to $\hat{\sigma}$. Zhang et al. [2] showed that the minimax convergence of mismatch rate as $\frac{nI}{K \log K} \to \infty$ is

$$
\inf_{\hat{\sigma}} \sup_{\Theta(n,K,\theta_{in},\theta_{out},\beta)} \mathbb{E} r(\sigma, \hat{\sigma}) = \begin{cases} \exp\left\{-(1 + o(1))\frac{nI}{2}\right\}, & K = 2, \\ \exp\left\{-(1 + o(1))\frac{nI}{\beta K}\right\}, & K \geq 3, \end{cases} \tag{3}
$$

where $1 + \epsilon_n \leq \beta < \sqrt{5/3}$ for some $\epsilon_n = CK/n$ with a large enough constant $C$. Here, $I$ is a constant related to minimum in-cluster and maximum out-of-cluster connectivity probabilities $\theta_{in}$ and $\theta_{out}$ and is defined as

$$
I := -2 \log\Big(\sqrt{\theta_{in}\theta_{out}} + \sqrt{(1 - \theta_{in})(1 - \theta_{out})}\Big). \tag{4}
$$

Furthermore, Zhang et al. [2] showed that $I$ is asymptotically equivalent to

$$\tilde{I} := \frac{(\theta_{in} - \theta_{out})^2}{\theta_{in}}. \tag{5}$$

$\tilde{I}$ can be interpreted as the "signal-to-noise ratio" of the graph. A larger $\tilde{I}$, hence $I$, leads to a faster convergence of mismatch rate to 0. We utilize this notion of "signal-to-noise ratio" to investigate the impact of aggregation. Specifically, given adjacency matrices $A_1$ and $A_2$ of two independent SBM graphs on the same set of cells, we use superscript to indicate their respective parameters, e.g., $\theta_{k\ell}^{A_1}$ represents the connectivity probability of cells in cluster $k$ and cells in cluster $\ell$ in graph $A_1$. Next, we define the adjacency matrix of "union graph" $B$ as

$$B_{i,j} := \begin{cases} 1, & \text{if } (A_1)_{ij} = 1 \text{ or } (A_2)_{ij} = 1, \\ 0, & \text{o.w.,} \end{cases} \tag{6}$$

where $(A_1)_{ij}$ denotes the entry at the $i$-th row and $j$-th column of the adjacency matrix $A_1$. Then, $\theta_{k\ell}^{B}$, the connectivity probability of cells in cluster $k$ and cells in cluster $\ell$ in the union graph is given by

$$\begin{aligned} \theta_{k\ell}^{B} :=& \mathbb{P}\Big(B_{i,j} = 1 \Big| \sigma(i) = k, \sigma(j) = \ell\Big) \\ =& 1 - \mathbb{P}\Big((A_1)_{i,j} = 0 \text{ and } (A_2)_{i,j} = 0 \Big| \sigma(i) = k, \sigma(j) = \ell\Big) \\ =& 1 - \mathbb{P}\Big((A_1)_{i,j} = 0 \Big| \sigma(i) = k, \sigma(j) = \ell\Big) \times \\ & \mathbb{P}\Big((A_2)_{i,j} = 0 \Big| \sigma(i) = k, \sigma(j) = \ell\Big) \\ =& 1 - (1 - \theta_{k\ell}^{A_1})(1 - \theta_{k\ell}^{A_2}) \\ =& \theta_{k\ell}^{A_1} + \theta_{k\ell}^{A_2} - \theta_{k\ell}^{A_1}\theta_{k\ell}^{A_2}. \end{aligned} \tag{7}$$

The resulting union graph has the stochastic block structure with minimum in-cluster connectivity $\theta_{in}^{B} \geq \theta_{in}^{A_1} + \theta_{in}^{A_2} - \theta_{in}^{A_1}\theta_{in}^{A_2}$, and maximum out-of-cluster connectivity $\theta_{out}^{B} \leq \theta_{out}^{A_1} + \theta_{out}^{A_2} - \theta_{out}^{A_1}\theta_{out}^{A_2}$.

Let $\tilde{I}^{A_1}$, $\tilde{I}^{A_2}$ and $\tilde{I}^{B}$ denote the signal-to-noise ratios of graphs $A_1$, $A_2$, and $B$, respectively.

Then, we have

$$\tilde{I}^B = \frac{(\theta_{in}^B - \theta_{out}^B)^2}{\theta_{in}^B} \geq \frac{(\theta_{in}^{A_1} + \theta_{in}^{A_2} - \theta_{in}^{A_1}\theta_{in}^{A_2} - \theta_{out}^{A_1} - \theta_{out}^{A_2} + \theta_{out}^{A_1}\theta_{out}^{A_2})^2}{\theta_{in}^{A_1} + \theta_{in}^{A_2} - \theta_{in}^{A_1}\theta_{in}^{A_2}}. \tag{8}$$

Under $\theta_{in}^{A_1} = \theta_{in}^{A_2}$ and $\theta_{out}^{A_1} = \theta_{out}^{A_2}$, we have

$$\tilde{I}^B \geq \frac{(2 - \theta_{in}^{A_1} - \theta_{out}^{A_1})^2}{2 - \theta_{in}^{A_1}} \times \frac{(\theta_{in}^{A_1} - \theta_{out}^{A_1})^2}{\theta_{in}^{A_1}} = \tilde{I}_{A_1} \times \frac{(2 - \theta_{in}^{A_1} - \theta_{out}^{A_1})^2}{2 - \theta_{in}^{A_1}}. \tag{9}$$

This relation implies that when $A_1$, $A_2$ are very sparse, i.e., with small $\theta_{in}^{A_1}$ and $\theta_{out}^{A_1}$, the signal-to-noise ratio $\tilde{I}_B$ of the union graph is almost twice as large as the individual graphs $\tilde{I}_{A_1}$ and $\tilde{I}_{A_2}$.

### Dimension reduction perturbs graph neighbors

We analyzed the general impact of dimension reduction on the local neighbourhoods of the cells under a simple model where data for each cell is from a mixture of Gaussian distributions with $K+1$ components. Specifically, let $y_0$, $y_i$, and $y_j$ denote the expression vectors of three cells drawn from the Gaussian mixture $\sum_{k=0}^{K} p_k \mathcal{N}(\mu_k, \sigma^2 I_d)$, where $\mathcal{N}(.,.)$ denotes the Gaussian probability density function. Here, $K+1$ is the number of clusters, i.e., cell types, $d$ is the dimension of the data from the Gaussian mixture, i.e., numbers of genes, $p_k$'s with $\sum_{k=0}^{K} p_k = 1$ are the mixing probabilities of the mixture distribution and represent the population proportion of each cell type. Without loss of generality, suppose a cell indexed by 0 and with expression profile $y_0$ is from cluster 0, and cells $i$ and $j$ with expression profiles $y_i, y_j$ are from cluster 1. Next, we consider projecting the $d$-dimensional ($d$ very large) $y_0$, $y_i$, and $y_j$ onto a $d_0$-dimensional space ($d_0$ very small) and investigate the relative ordering of the distances between cells $i$, $j$ to cell 0 in the $d_0$-dimensional space to their orderings in the $d$-dimensional space.

The following lemma shows diminishing dependence between these two sets of orderings which implies low levels of edge overlap between the shared nearest neighbor graph in the low-dimensional space and the shared nearest neighbor graph constructed from the original high dimensional data.

**Lemma 0.1.** *Suppose* $y_i, y_j \in \mathcal{N}(\mu_1, \sigma^2 I_d)$*,* $y_0 \in \mathcal{N}(\mu_0, \sigma^2 I_d)$*,* $Z = [z_1, \ldots, z_{d_0}] \in \mathbb{R}^{d\times d_0}$*, where* $\{z_t\}_{t=1,\ldots,d_0}$ *are linearly independent unit vectors. Define* $u_t = (Z^T Z)^{-1} Z^T y_t$ *for* $t = 0, i, j$*. If* $d_0$ *is fixed,* $\|\mu_1 - \mu_0\|_4$ *is bounded as* $d \to \infty$*, then we have*

$$\Big| \mathbb{P}\left[\|y_i - y_0\|^2 > \|y_j - y_0\|^2, \|u_i - u_0\|^2 > \|u_j - u_0\|^2\right] - \mathbb{P}\left[\|y_i - y_0\|^2 > \|y_j - y_0\|^2\right] \mathbb{P}\left[\|u_i - u_0\|^2 > \|u_j - u_0\|^2\right] \Big| \to 0$$

*Proof.* Define $H = Z(Z^T Z)^{-1} Z^T$ as the projection matrix onto $d_0$ dimensional space, and the orthogonal complement of $H$ as $H^\perp = I - H$.

$$\begin{aligned} \mathbb{E}\Big|\|H(y_i - y_0)\|^2 - \|H(y_j - y_0)\|^2\Big| &\leq 2\mathbb{E}\|H(y_i - y_0)\|^2 & (10) \\ &= 4d_0\sigma^2 + 2(\mu_1 - \mu_0)^T H(\mu_1 - \mu_0) & (11) \\ &\leq 4d_0\sigma^2 + 2\|\mu_1 - \mu_0\|^2. & (12) \end{aligned}$$

By Markov's inequality, for any $t > 0$,

$$\mathbb{P}\Big\{\Big|\|H(y_i - y_0)\|^2 - \|H(y_j - y_0)\|^2\Big| \leq t\Big\} \geq 1 - \frac{4d_0\sigma^2 + 2\|\mu_1 - \mu_0\|^2}{t}. \quad (13)$$

Suppose the eigenvalue decomposition of projection matrix $H^\perp$ has the form $H^\perp = U I_{d-d_0} U^T$. Define $v_t = U^T(y_t - y_0)$, for $t = i, j$, and $\tilde{\mu} = U^T(\mu_1 - \mu_0)$. Then $v_{tk} \sim \mathcal{N}(\tilde{\mu}_k, 2\sigma^2)$ and $v_{tk}$ are mutually independent for $t = i, j$ and $k = 1, \ldots, d - d_0$, and

$$\|H^\perp(y_i - y_0)\|^2 - \|H^\perp(y_j - y_0)\|^2 = \sum_{k=0}^{d-d_0} (v_{i,k}^2 - v_{jk}^2), \quad (14)$$

$$\mathbb{E}\left[\frac{v_{ik}^2 - v_{jk}^2}{\sigma^2}\right] = 0, \ \mathrm{Var}\left(\frac{v_{ik}^2 - v_{jk}^2}{\sigma^2}\right) = 64 + 32\Big(\frac{\tilde{\mu}_k}{\sigma}\Big)^2. \quad (15)$$

Taking $s_d^2 = 32\sum_{k=1}^{d-d_0}[2 + \left(\frac{\tilde{\mu}_k}{\sigma}\right)^2] = 64(d - d_0) + 32\frac{(\mu_1-\mu_0)^T H^\perp(\mu_1-\mu_0)}{\sigma^2}$, we can verify the Feller's condition [3],

$$\lim_{d\to\infty}\sum_{k=1}^{d-d_0}\frac{\mathbb{E}[\frac{v_{ik}^2-v_{jk}^2}{\sigma^2}]^4}{\epsilon^2 s_d^4}=\frac{\sum_k 3072\big[(\frac{\tilde{\mu}_k}{\sigma})^4+12(\frac{\tilde{\mu}_k}{\sigma})^2+12\big]}{\epsilon^2\big[64(d-d_0)+32\frac{(\mu_1-y_0)^T H^\perp(\mu_1-y_0)}{\sigma^2}\big]^2}\to 0, \tag{16}$$

for any $\epsilon > 0$ and $d \to \infty$, and apply the Lindeberg central limit theorem [3] to get

$$\sum_k \frac{v_{ik}^2 - v_{jk}^2}{\sigma^2 s_d} \to^d \mathcal{N}(0,1). \tag{17}$$

This asymptotic result leads to the following probability bound

$$\begin{aligned}
&\mathbb{P}\Big(\Big|\|H^\perp(y_i-y_0)\|^2-\|H^\perp(y_j-y_0)\|^2\Big|>t\Big) && (18)\\
&\qquad=\mathbb{P}\Big(\frac{|\sum_{k=1}^{d-d_0}(v_{ik}^2-v_{jk}^2)|}{\sigma^2 s_d}>\frac{t}{\sigma^2 s_d}\Big) && (19)\\
&\qquad=2\left(1-\Phi\left(\frac{t}{\sigma^2 s_d}\right)\right)-o(1) && (20)\\
&\qquad=1-o(1), && (21)
\end{aligned}$$

as $d \to \infty$. Finally, we have

$$\begin{aligned}
\frac{1}{4} \quad &\geq \mathbb{P}\Big[\|y_i-y_0\|^2>\|y_j-y_0\|^2, \|u_i-u_0\|^2<\|u_j-u_0\|^2\Big] && (22)\\
&\geq \mathbb{P}\Big[\|H^\perp(y_i-y_0)\|^2-\|H^\perp(y_j-y_0)\|^2\geq t, && (23)\\
&\qquad \Big|\|H(y_i-y_0)\|^2-\|H(y_j-y_0)\|^2\Big|<t, \|u_i-u_0\|^2<\|u_j-u_0\|^2\Big]\\
&= \mathbb{P}\Big[\|H^\perp(y_i-y_0)\|^2-\|H^\perp(y_j-y_0)\|^2\geq t\Big] && (24)\\
&\qquad \times\mathbb{P}\Big[\Big|\|H(y_i-y_0)\|^2-\|H(y_j-y_0)\|^2\Big|<t, \|u_i-u_0\|^2<\|u_j-u_0\|^2\Big]\\
&\geq \left[\frac{1}{2}-o(1)\right]\times\left[\frac{1}{2}-o(1)\right] && (25)\\
&= \frac{1}{4}-o(1) && (26)
\end{aligned}$$

The last inequality holds with an appropriate choice of $t$. $\square$

### Supplementary figures

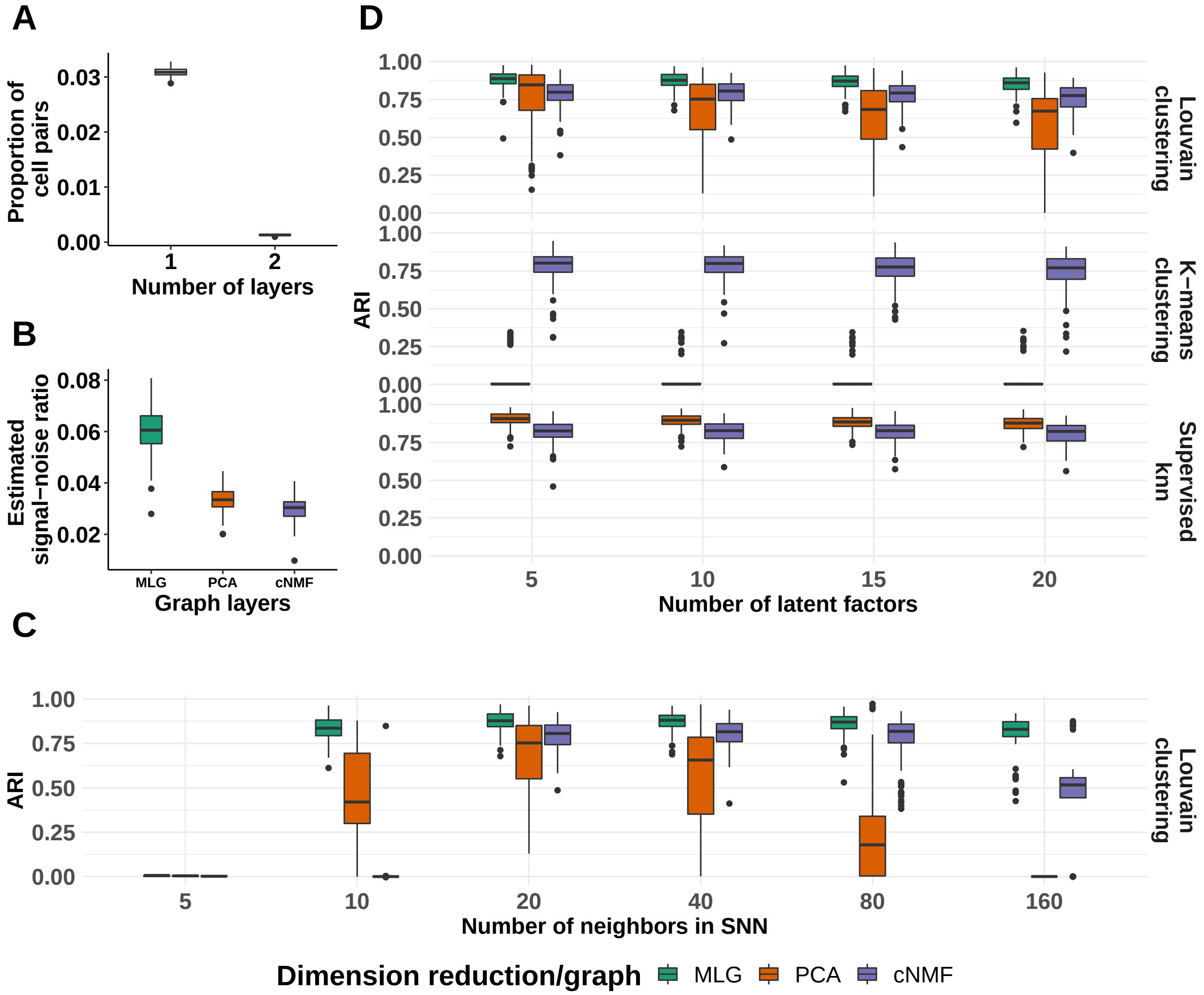


Figure S1: **Simulation results for the "small-condition-effect" setting.** (A) Proportions of cell pairs with edges across different numbers of layers of MLG constructed from SNN graphs of PCA and cNMF, Seurat, and Liger (with 20 neighbours in SNN graphs and 10 latent factors in low-dimensional embeddings). The boxplots depict the proportions across all the simulation replicates. (B) Estimated signal-to-noise ratios of SNN graphs constructed from different low-dimensional embeddings and their multilayer graph across all the simulation replicates (with 20 neighbours in SNNs and 10 latent factors in low-dimensional embeddings). (C) Louvain clustering accuracy of SNN graphs as a function of numbers of neighbors in SNN graph construction (with 10 latent factors in low-dimensional embeddings). (D) Adjusted Rand index comparison of Louvain and k-means clustering of SNN graphs from different low-dimensional embeddings and their MLG as a function of number of latent factors in the low-dimensional projections (with 20 neighbours in SNN graphs). ARI values of supervised knn classifiers for individual SNN graphs are provided as reference.

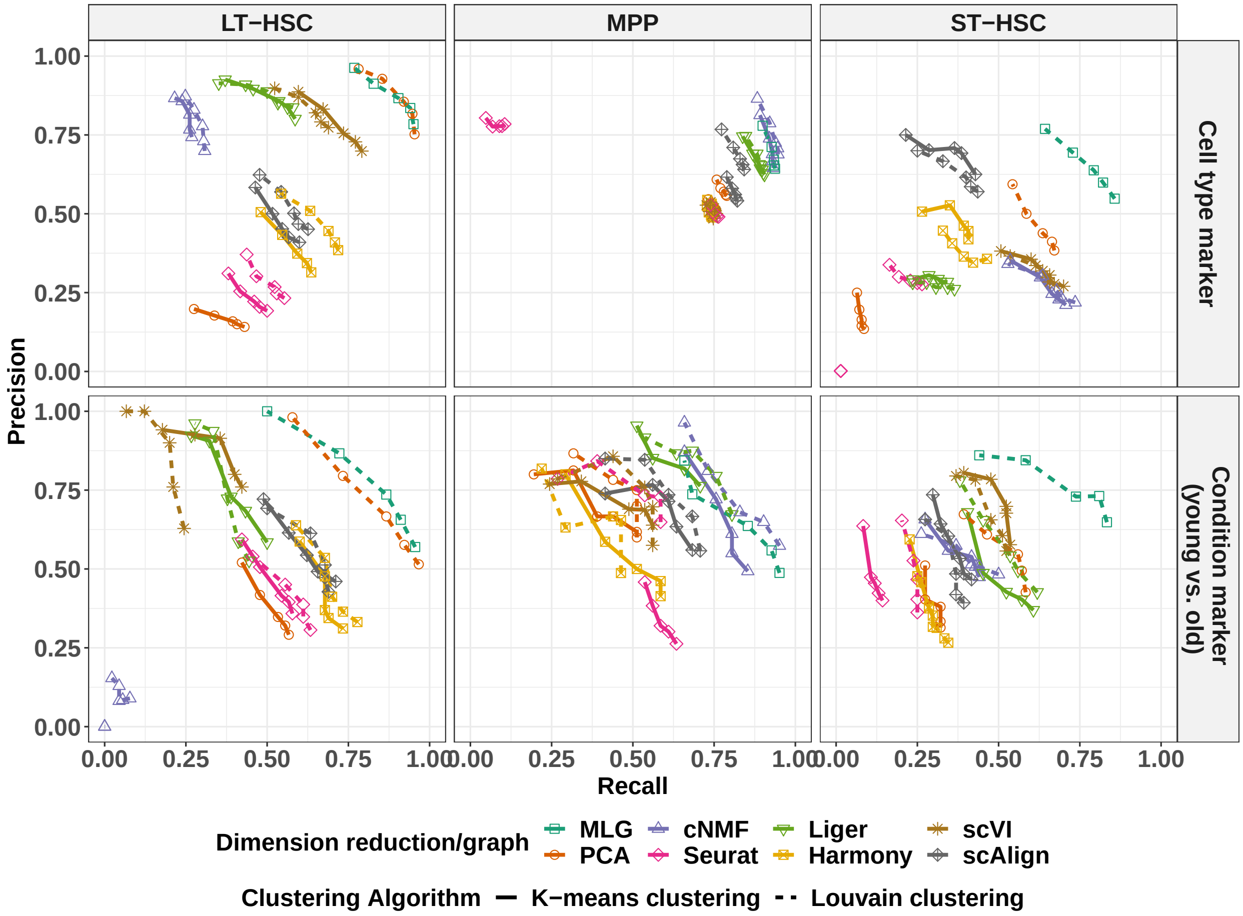


Figure S2: **Differential expression analysis for cell type/cluster marker gene and condition (young vs. old) DE gene identification in the *Kowalczyk_2* dataset.** The precision-recall (PR) values in the top panel evaluate inferred cluster (i.e., cell type) marker genes of each method against cell type marker genes defined with ground truth cell type labels as gold standard. The bottom panel evaluates the inferred condition (i.e.,age) DE genes of each cell type against the gold standard. Gold standard cell type marker and age DE genes are defined as genes with Bonferroni corrected p-values less than 0.01 in the differential expression analysis with ground truth cell labels. The PR values are reported at cutoffs of 0.2, 0.1, 0.05, 0.01, and 0.001 for Bonferroni adjusted p-values in the cluster marker gene and age DE gene identification analysis.

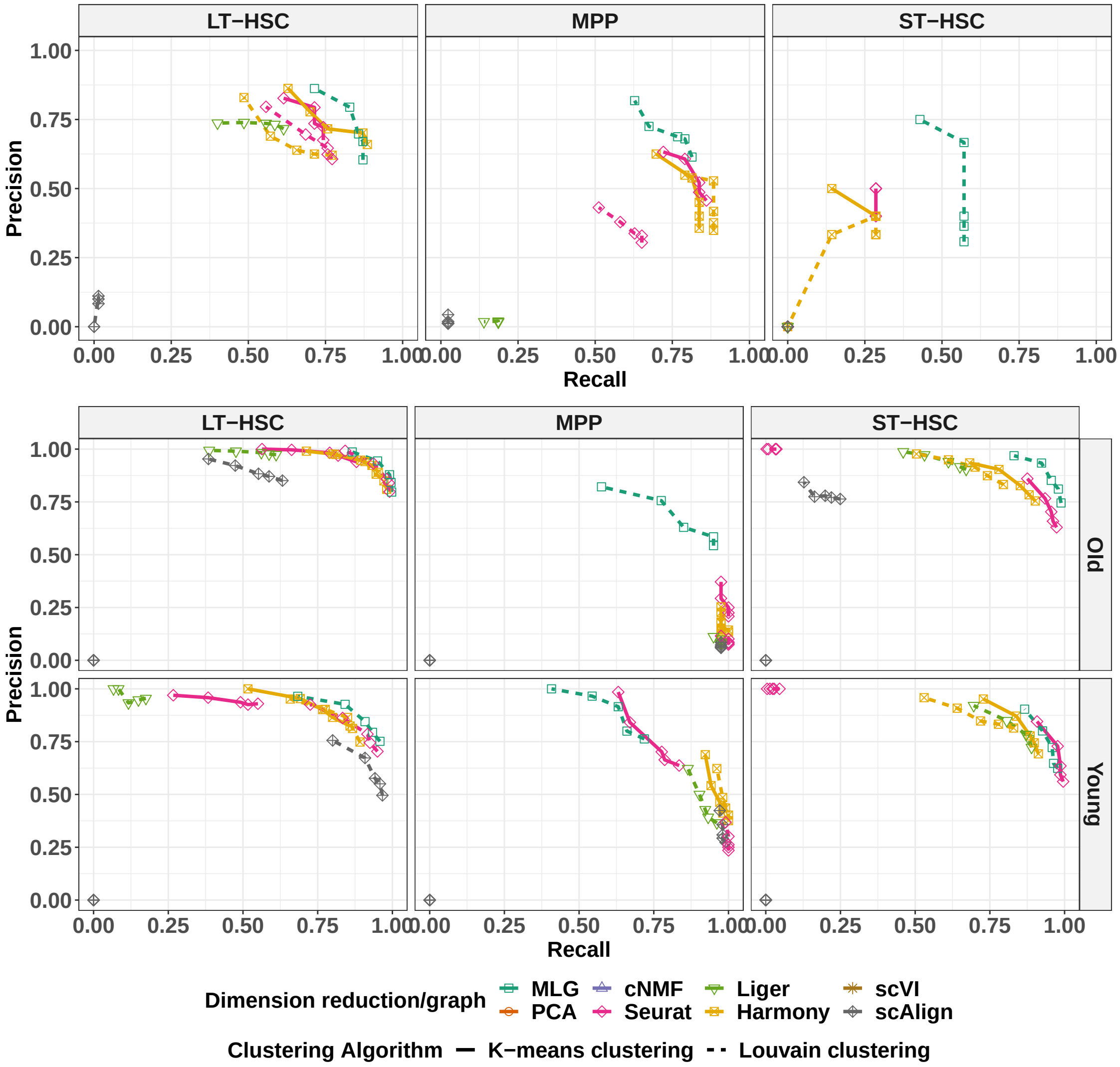


Figure S3: **Differential expression analysis for cell type/cluster marker gene and LPS+PAM stimuli DE gene identification in the *Mann* dataset.** Top panel precision-recall (PR) values evaluate inferred cluster, i.e., cell type, marker genes of each method with respect to cell type maker genes identified with ground truth cell type labels as gold standard. The bottom panel evaluates the inferred condition (i.e., LPS+PAM stimuli) DE genes of each cell type against the gold standard, separately for old and young mice. Gold standard cell type marker and LPS+PAM stimuli DE genes are defined as genes with Bonferroni corrected p-values less than 0.01 in the differential expression analysis with ground truth cell labels. The PR values are reported at cutoffs of 0.2, 0.1, 0.05, 0.01, 0.001 for Bonferroni adjusted p-values in the differential analysis for cluster marker genes and stimuli DE genes within each cluster. Several clustering of low-dimensional projections (k-means-PCA, Louvain-PCA, k-means-Liger, k-means-scVI, Louvain-scVI, k-means-cNMF, Louvain-cNMF) identified clusters with only cells from one condition, preventing a cell type-specific condition DE gene analysis.

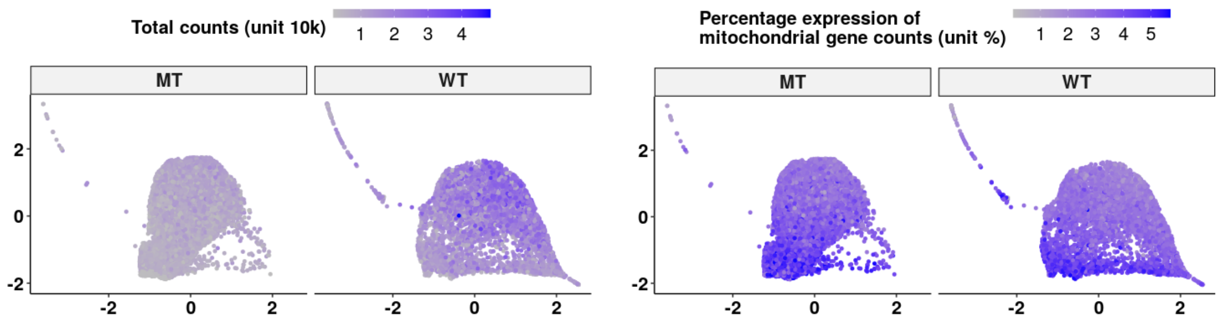


Figure S4: **SPRING visualization of the *Johnson_20* dataset, superimposed with total counts (left) and percentage expression of mitochondrial genes (right) per cell.**

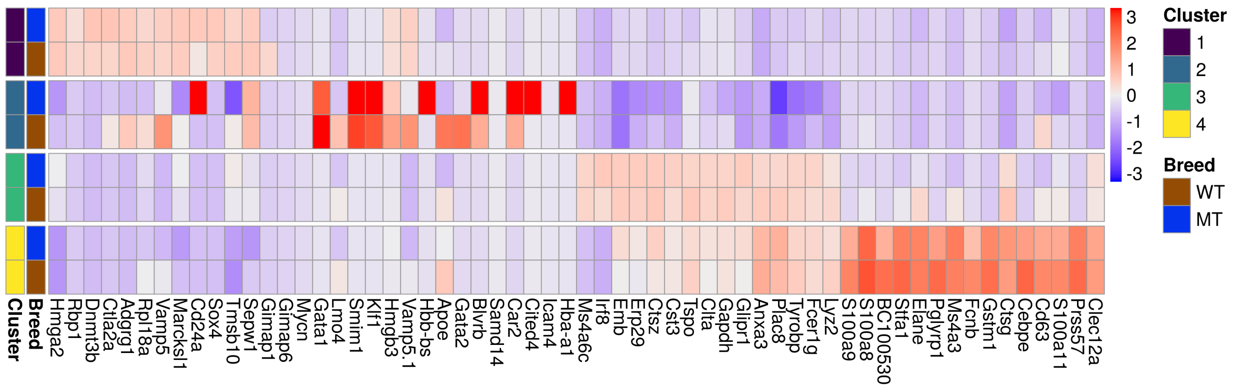


Figure S5: **Heatmap of median gene expression of MLG cluster marker genes, grouped by breed and MLG cluster, for the *Johnson_20* dataset.**

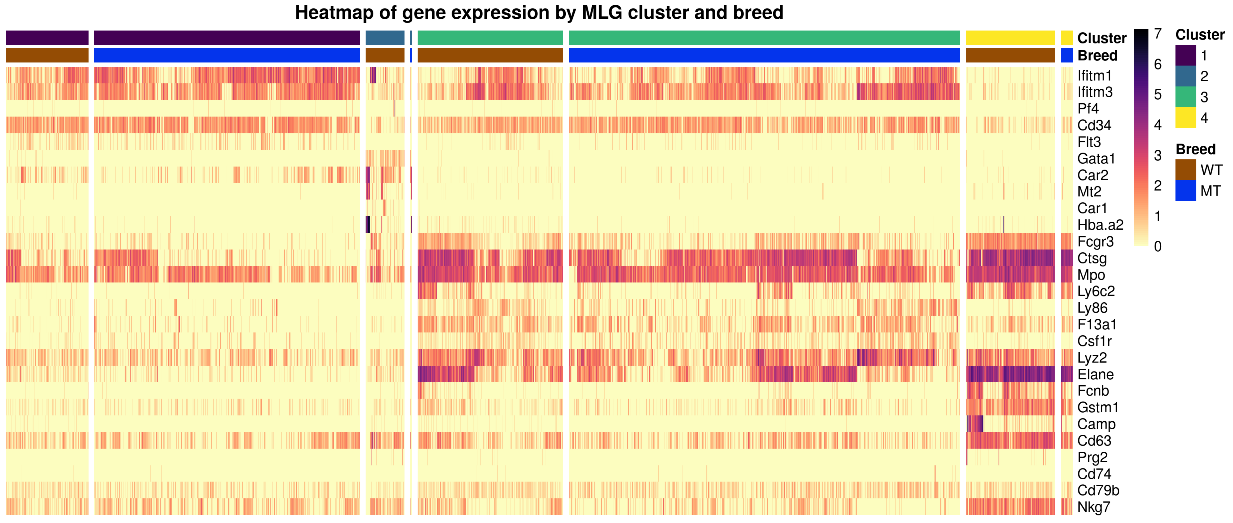


Figure S6: **Heatmap of cell-level gene expression of marker genes from [4], grouped by breed and MLG clustering of the *Johnson_20* dataset.**

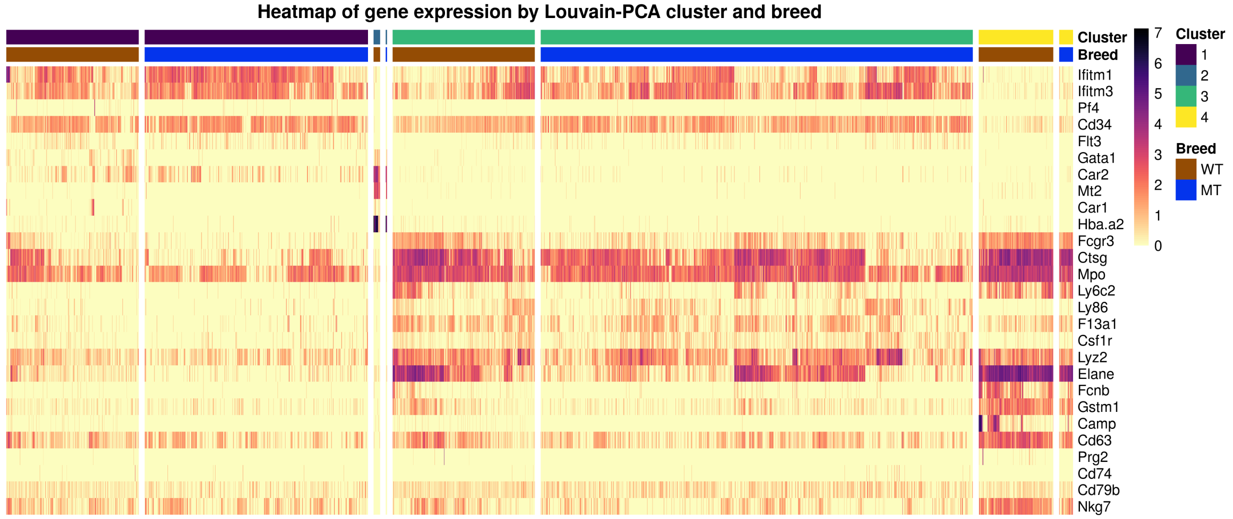


Figure S7: **Heatmap of cell-level gene expression of marker genes from [4], grouped by breed and Louvain clustering of PCA low-dimensional embedding of the *Johnson_20* dataset.**

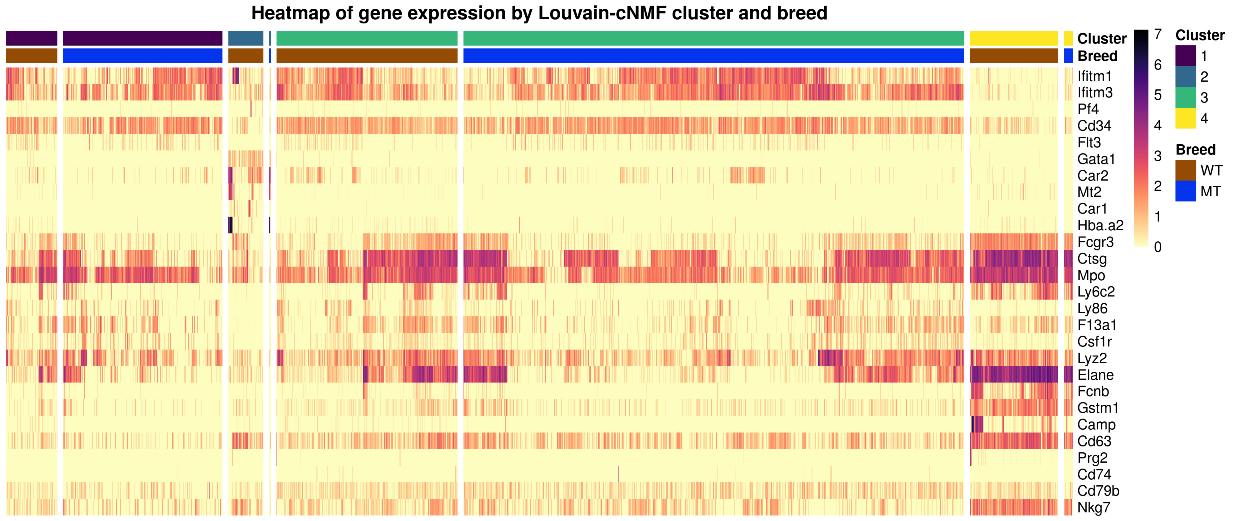


Figure S8: **Heatmap of cell-level gene expression of marker genes from [4], grouped by breed and Louvain clustering of cNMF low-dimensional embedding of the *Johnson_20* dataset.**

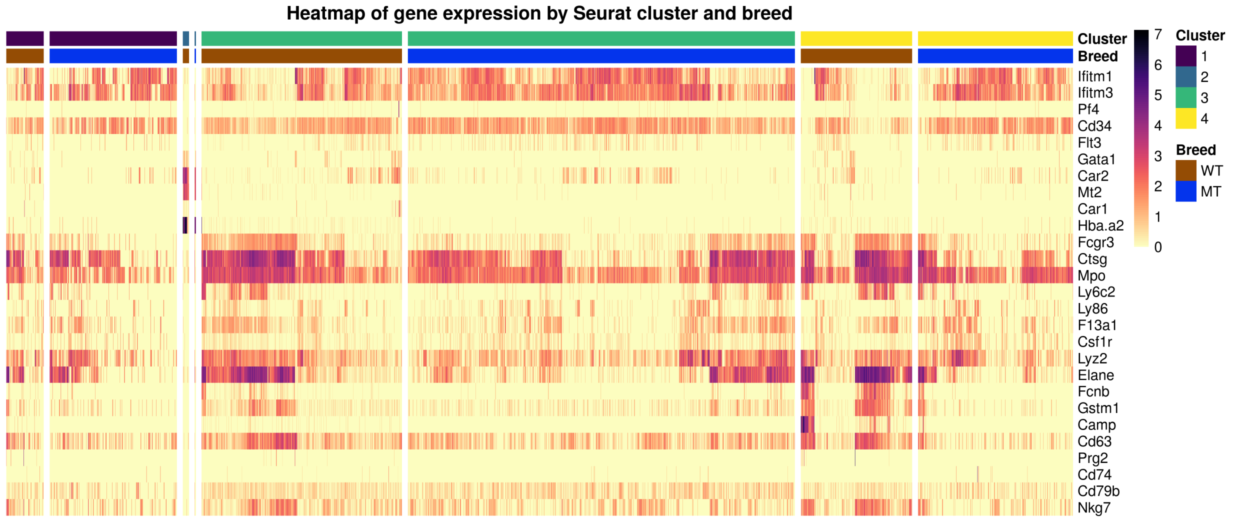


Figure S9: **Heatmap of cell-level gene expression of marker genes from [4], grouped by breed and Louvain clustering of Seurat-integration low-dimensional embedding of the *Johnson_20* dataset.**

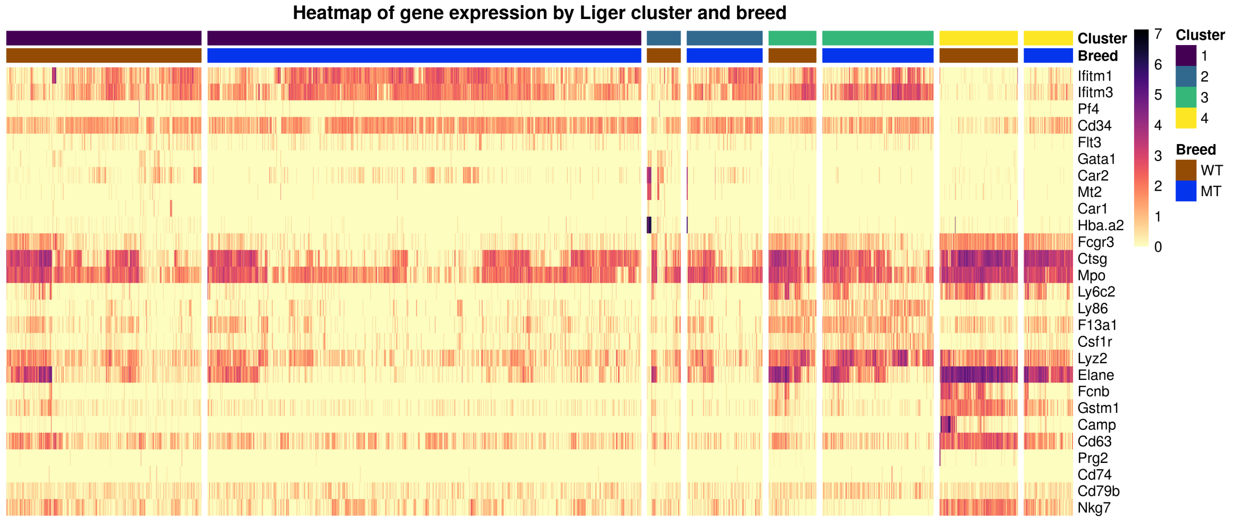


Figure S10: **Heatmap of cell-level gene expression of marker genes from [4], grouped by breed and Louvain clustering of Liger low-dimensional embedding of the *Johnson_20* dataset.**

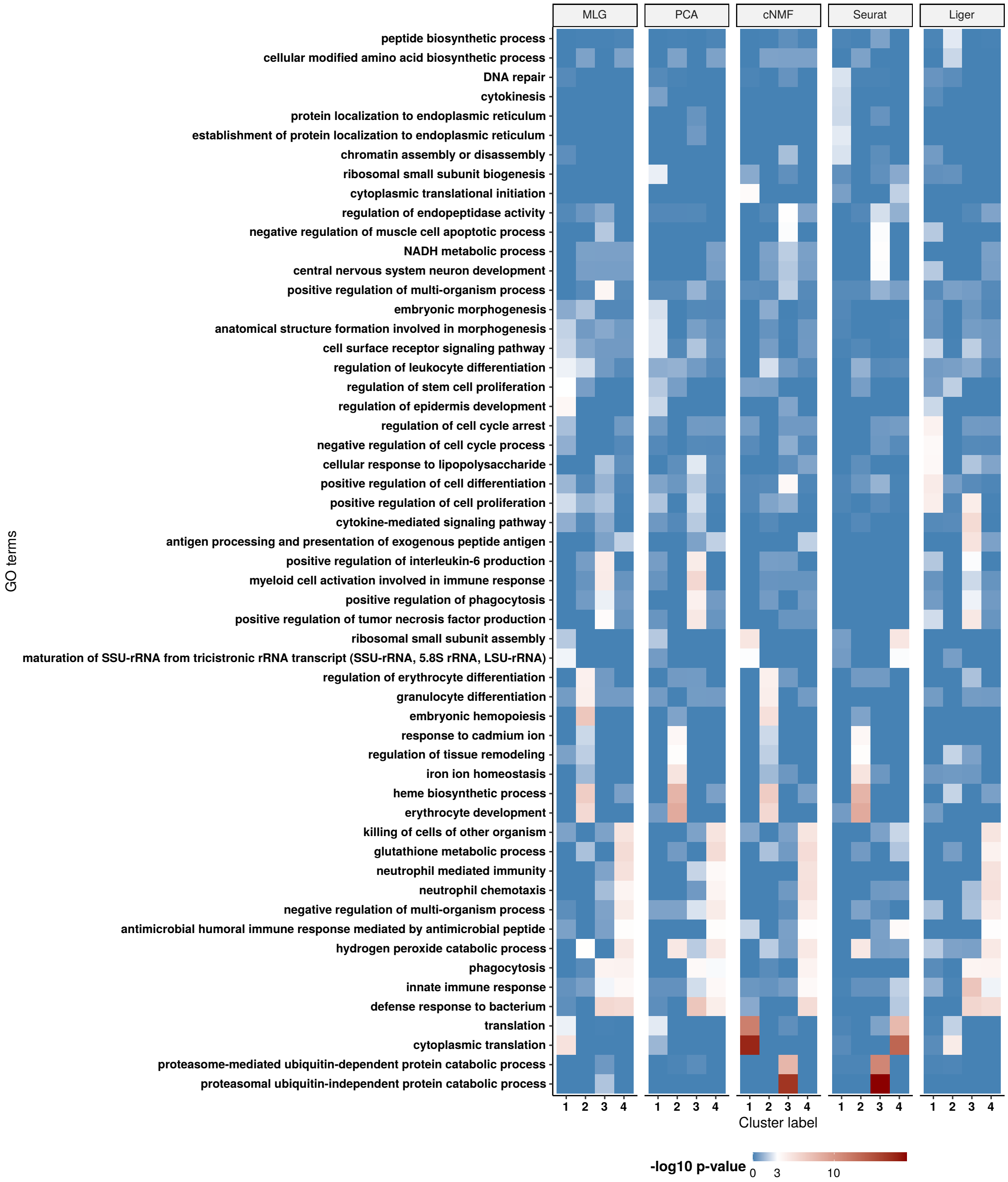


Figure S11: **Gene set enrichment analysis for the *Johnson_20* dataset.** Heatmap of the -$\log_{10}$ enrichment p-values of the GO terms across clusters identified by different methos. Top 5 GO terms from gene set enrichment analysis of each cluster of each individual method are pooled together.

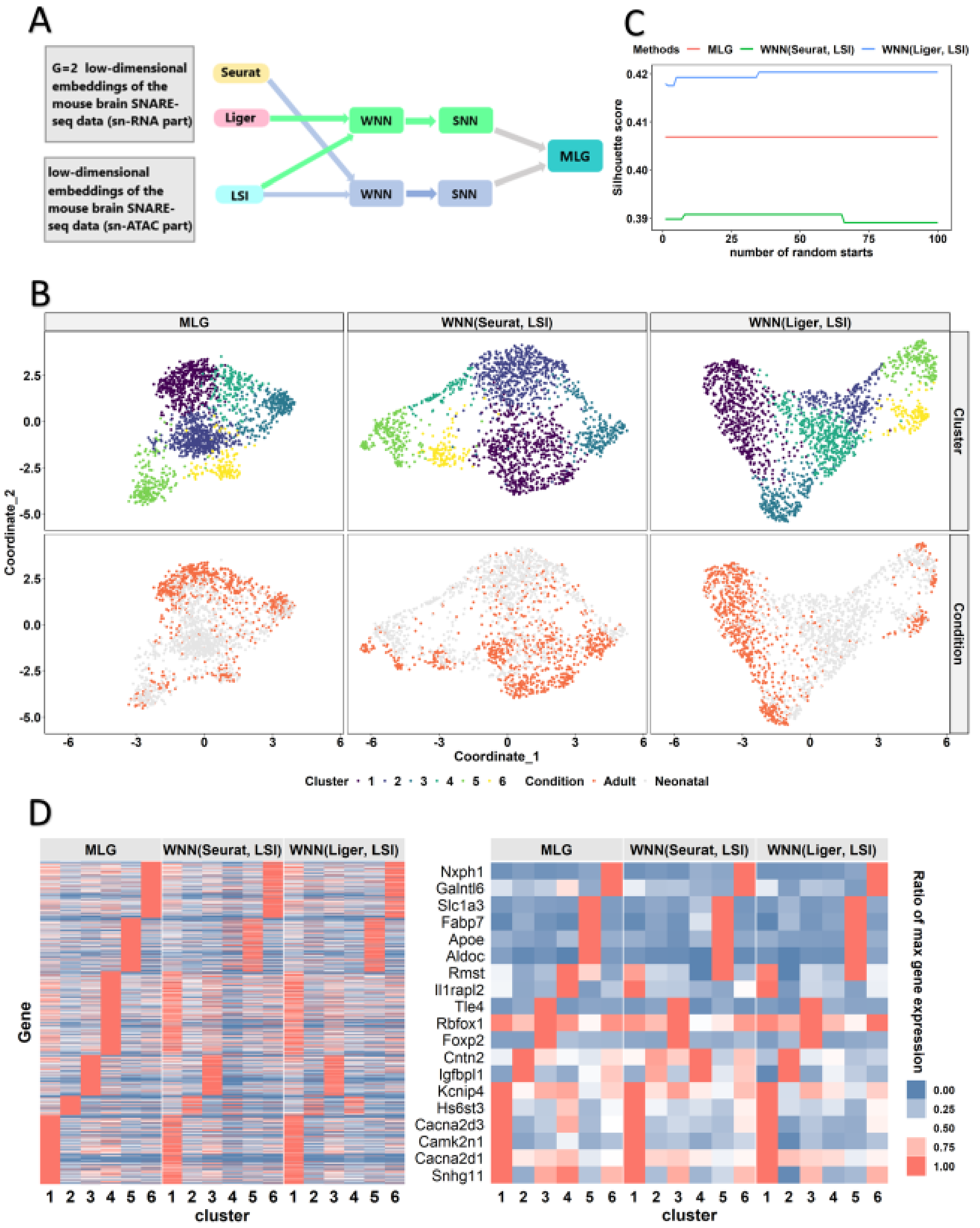


Figure S12: **Application to dataset *Chen19* (replicate 2).** (A) Data processing workflow of MLG for SNARE-seq. Dimension reduction methods Liger and Seurat are applied to snRNA-seq, Latent Semantic Indexing (LSI) is applied to snATAC-Seq. Weighted nearest neighbor (WNN) graphs [5] are constructed using Liger-LSI and Seurat-LSI low-dimensional embeddings. Finally, the two WNN graphs are inputted to the MLG framework. (B) Visualization of clusters and conditions on the force-directed layout of MLG and the UMAP coordinates of WNN(Seurat, LSI) and WNN(Liger, LSI). (C) Average silhouette scores for the three clustering methods, computed with the corresponding UMAP or force-directed layout coordinates. X-axis is the number of random starts inputted to the Louvain algorithm. (D) Heatmap of the scaled gene expression for all the genes (left) and selected genes (on the right).

**Supplementary tables**

| Dataset | # of cells | System | Conditions/Stimuli |
|---|---|---|---|
| *Kowalczyk_1* [6] | 1,058 | HSPC | Young (2-3 months) and old (22 months) mice. |
| *Kowalczyk_2* [6] | 1,428 | HSPC | Young (2-3 months) and old (22 months) mice. |
| *Mann* [7] | 949 | HSPC | Young (8-12 weeks)<br>and old (20-24 months) mice<br>with and without LPS+PAM stimulation. |
| *Cellbench* [8] | 636 | Synthetic cells | Two sequencing protocols:<br>CEL-seq-2 and SORT-seq. |

Table S1: Benchmark datasets.

Setting 1

| | Cell population proportion | | | Activity GEP 1 usage | | | Activity GEP 2 usage | | |
|---|---|---|---|---|---|---|---|---|---|
| cell type | 1 | 2 | 3 | 1 | 2 | 3 | 1 | 2 | 3 |
| condition 1 | 12.5% | 25% | 12.5% | 0 | 0 | 5% | 40% | 40% | 40% |
| condition 2 | 15% | 15% | 20% | 25% | 20% | 20% | 5% | 0 | 0 |

Setting 2

| | Cell population proportion | | | Activity GEP 1 usage | | | Activity GEP 2 usage | | |
|---|---|---|---|---|---|---|---|---|---|
| cell type | 1 | 2 | 3 | 1 | 2 | 3 | 1 | 2 | 3 |
| condition 1 | 12.5% | 25% | 12.5% | 5% | 10% | 5% | 30% | 25% | 20% |
| condition 2 | 15% | 15% | 20% | 30% | 25% | 25% | 5% | 0 | 10% |

Table S2: Simulation settings 1 and 2 correspond to the large and small condition effects, respectively.

[3] Rick Durrett. *Probability: theory and examples*, volume 49. Cambridge university press, 2019.

[4] Amir Giladi, Franziska Paul, Yoni Herzog, Yaniv Lubling, Assaf Weiner, Ido Yofe, Diego Jaitin, Nina Cabezas-Wallscheid, Regine Dress, Florent Ginhoux, et al. Single-cell characterization of

haematopoietic progenitors and their trajectories in homeostasis and perturbed haematopoiesis. *Nature cell biology*, 20(7):836–846, 2018.

[5] Yuhan Hao, Stephanie Hao, Erica Andersen-Nissen, William M Mauck, Shiwei Zheng, Andrew Butler, Maddie Jane Lee, Aaron J Wilk, Charlotte Darby, Michael Zagar, et al. Integrated analysis of multimodal single-cell data. *bioRxiv*, 2020.

[6] Monika S Kowalczyk, Itay Tirosh, Dirk Heckl, Tata Nageswara Rao, Atray Dixit, Brian J Haas, Rebekka K Schneider, Amy J Wagers, Benjamin L Ebert, and Aviv Regev. Single-cell rna-seq reveals changes in cell cycle and differentiation programs upon aging of hematopoietic stem cells. *Genome research*, 25(12):1860–1872, 2015.

[7] Mati Mann, Arnav Mehta, Carl G de Boer, Monika S Kowalczyk, Kevin Lee, Pearce Haldeman, Noga Rogel, Abigail R Knecht, Daneyal Farouq, Aviv Regev, et al. Heterogeneous responses of hematopoietic stem cells to inflammatory stimuli are altered with age. *Cell reports*, 25(11): 2992–3005, 2018.

[8] Luyi Tian, Xueyi Dong, Saskia Freytag, Kim-Anh Le Cao, Shian Su, Abolfazl JalalAbadi, Daniela Amann-Zalcenstein, Tom S Weber, Azadeh Seidi, Jafar S Jabbari, et al. Benchmarking single cell rna-sequencing analysis pipelines using mixture control experiments. *Nature methods*, 16(6):479–487, 2019.
